# Revision of splicing variants in the *DMD* gene

**DOI:** 10.1101/2024.01.31.578175

**Authors:** Kseniya Davydenko, Alexandra Filatova, Mikhail Skoblov

**Affiliations:** Research Centre of Medical Genetics, Laboratory of Functional Genomics, 115478 Moscow, Russia

**Keywords:** *DMD*, Duchenne muscular dystrophy, Becker muscular dystrophy, splicing variants, minigene

## Abstract

**Background:** Pathogenic variants in the dystrophin (*DMD*) gene lead to X-linked recessive Duchenne muscular dystrophy (DMD) and Becker muscular dystrophy (BMD). Nucleotide variants that affect splicing are a known cause of hereditary diseases. However, their representation in the public genomic variation databases is limited due to the low accuracy of their interpretation, especially if they are located within exons. The analysis of splicing variants in the *DMD* gene is essential both for understanding the underlying molecular mechanisms of the dystrophinopathies’ pathogenesis and selecting suitable therapies for patients.

**Results:** Using deep *in silico* mutagenesis of the entire *DMD* gene sequence and subsequent SpliceAI splicing predictions, we identified 7,948 *DMD* single nucleotide variants that could potentially affect splicing, 863 of them were located in exons. Next, we analyzed over 1,300 disease-associated *DMD* SNVs previously reported in the literature (373 exonic and 956 intronic) and intersected them with SpliceAI predictions. We predicted that ∼95% of the intronic and ∼10% of the exonic reported variants could actually affect splicing. Interestingly, the majority (75%) of patient-derived intronic variants were located in the AG-GT terminal dinucleotides of the introns, while these positions accounted for only 13% of all intronic variants predicted *in silico*. Of the 97 potentially spliceogenic exonic variants previously reported in patients with dystrophinopathy, we selected 38 for experimental validation. For this, we developed and tested a minigene expression system encompassing 27 *DMD* exons. The results showed that 35 (19 missense, 9 synonymous, and 7 nonsense) of the 38 *DMD* exonic variants tested actually disrupted splicing. We compared the observed consequences of splicing changes between variants leading to severe Duchenne and milder Becker muscular dystrophy and showed a significant difference in their distribution. This finding provides extended insights into relations between molecular consequences of splicing variants and the clinical features.

**Conclusions:** Our comprehensive bioinformatics analysis, combined with experimental validation, improves the interpretation of splicing variants in the *DMD* gene. The new insights into the molecular mechanisms of pathogenicity of exonic single nucleotide variants contribute to a better understanding of the clinical features observed in patients with Duchenne and Becker muscular dystrophy.

## Introduction

The *DMD* gene, located on the X chromosome, encodes dystrophin, an inner plasma membrane protein found in muscle fibers. Dystrophin consists of four distinct domains: the N-terminal actin-binding domain (ABD1), a central rod domain with 24 spectrin-like repeats and a second actin-binding domain (ABD2), a cysteine-rich domain (CRD), and the C-terminal domain (CTD) (1). Dystrophin plays a crucial role in connecting the sarcolemma and actin of the cytoskeleton, acting as a “shock absorber” during muscle contraction (2).

Dysfunction of dystrophin caused by pathogenic variants in the *DMD* gene leads to various conditions, including Duchenne muscular dystrophy (DMD), Becker muscular dystrophy (BMD), intermediate muscular dystrophy (IMD), and pure cardiac X-linked dilated cardiomyopathy (XLCM) (3, 4). The severity of these conditions is associated with the “reading frame rule,” which determines whether semi-functional dystrophin with preserved N-terminal and C-terminal protein-binding domains can be expressed (3). DMD is a severe form of progressive muscular atrophy that affects approximately one in 6000 live male births (4). It is caused by pathogenic variants that disrupt dystrophin synthesis, either by shifting the reading frame or generating a premature stop codon, resulting in a loss of protein function. BMD is a milder form of dystrophinopathy typically caused by pathogenic non-frameshift variants, which allow the expression of a shortened but partially functional dystrophin, leading to a later onset and slower disease progression (5).

The *DMD* gene is one of the largest genes in the human genome, spanning 2.3 Mb (6). Its substantial size and complicated structure contribute to a relatively high rate of spontaneous mutations, and it has been estimated that one in three cases of DMD caused by *de novo* substitutions. (7). The majority of pathogenic variants in the *DMD* gene (70–80%) are large rearrangements, including deletions and duplications affecting one or more exons (8). The remaining 20-30% of pathogenic variants include small rearrangements and single nucleotide substitutions. Single nucleotide variants (SNVs) that affect splicing account for 12–26% of short variants or 3–6% of all pathogenic variants in the *DMD* gene (9–11).

While single nucleotide substitutions in the *DMD* gene have been detected through targeted gene panel testing and whole exome sequencing, these methods primarily focus on coding regions, with limited coverage of intronic sequences outside canonical splice sites. Therefore, such approaches will miss splicing variants located deeper in introns (12). Moreover, splicing-affecting variants can sometimes be located in exons, disguising themselves as synonymous, missense, and even nonsense variants. Estimates suggest that splicing variants account for 10 to 30% of all pathogenic variants across various diseases (13–15). Several bioinformatics approaches exist for predicting the potential effects of SNVs on splicing during clinical genetic testing for hereditary diseases (16, 17). However, accurate identification of such variants and understanding their precise impact on splicing require experimental approaches.

Various experimental approaches are available to assess the pathogenicity of splicing-affecting variants. If a variant is present in a gene expressed in clinically accessible tissues (e.g., blood, skin, buccal epithelium), mRNA structure analysis through reverse transcription-polymerase chain reaction (RT-PCR) can be employed. For genes with low expression levels in available tissues, a minigene assay is used (18). A minigene consists of a minimal fragment of the target gene, such as a specific exon with surrounding intron sequences, cloned into an expression plasmid. This approach allows for the analysis of pre-mRNA splicing, confirming whether the variant affects splicing efficiency or activates alternative splicing sites. It also helps uncover the role of cis-acting elements in splicing regulation (19).

In this study, we performed a comprehensive analysis to assess the full spectrum of splicing variants of the *DMD* gene. We developed a minigene expression system to functionally evaluate the impact of exonic SNVs on pre-mRNA splicing and determined the contribution of these variants to the overall composition of pathogenic variants in the *DMD* gene. These findings elucidated the molecular mechanism of pathogenicity associated with exonic splicing variants and shed light on the relationship between genotype and disease severity. Accurate molecular diagnosis is crucial for patients with dystrophinopathy, primarily for optimal care and family planning. Furthermore, understanding the underlying pathogenic mechanisms becomes increasingly important in the context of actively developed therapeutic approaches, such as stop-codon readthrough and antisense oligonucleotide (AON)-mediated exon-skipping therapy (20–22).

## Results

### In silico analysis of all possible splicing SNVs in the DMD gene

We conducted an *in silico* analysis to estimate the number of potentially spliceogenic variants in the *DMD* gene. The entire gene sequence was subjected to deep mutagenesis, where each nucleotide was substituted with three other nucleotides. To assess the effect of these variants on splicing, we utilized the SpliceAI tool (23), which provides a Delta score (DS) ranging from 0 to 1. A higher DS indicates a greater probability of the variant being splice-altering.

Based on SpliceAI’s DS threshold of >0.2 (high recall/likely pathogenic), we identified 7,948 *DMD* variants likely to affect splicing (Table S2). The majority of these variants (89%) were found in intronic regions (Figure 1-A, 2). Among them, only 13% were located at the canonical AG-GT terminal dinucleotides of the intron, while an additional 13% were present in other nucleotides within the donor and acceptor splicing sites. Notably, most of the predicted intronic variants (71%) were located in deep intronic regions (>50 nt from exon-intron boundaries) (Figure 1-С).

**Figure 1.**
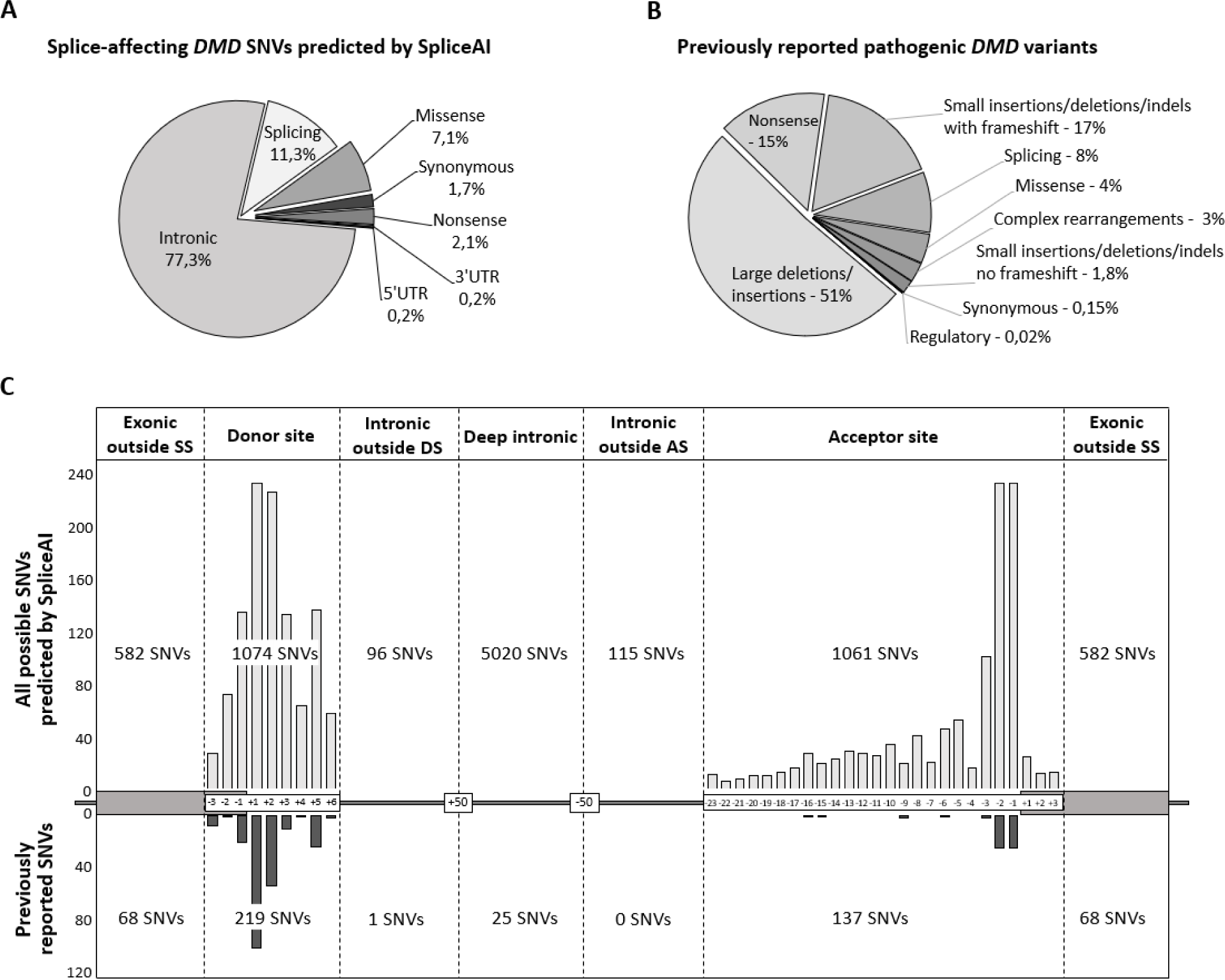
A. Distribution of pathogenic variants in the *DMD* gene reported in the scientific literature. B. Distribution of all possible single nucleotide variants potentially affecting splicing in the *DMD* gene according to the SpliceAI prediction. C. Number and localization of all possible spliceogenic *DMD* SNVs predicted by SpliceAI (top) and previously reported pathogenic variants in the *DMD* gene that could affect splicing according to the SpliceAI prediction (bottom). *Abbreviations: SS, Splicing sites; DS, Donor site; AS, Acceptor site*.

The annotation of exonic variants is typically based on the resulting amino acid sequence changes. In the case of nonsense variants, the molecular mechanism of pathogenicity is usually evident, while the impact of synonymous and missense variants may remain unclear, particularly if they occur outside of functional domains. However, it is known that some exonic variants can disrupt splicing, by either creating new splicing sites or disrupting existing ones. In such cases, the pathogenicity mechanism is primarily associated with mRNA structure or expression disruption, and these variants may lead to more severe or milder consequences than corresponding single amino acid substitutions. Using *in silico* mutagenesis approach, we identified 863 exonic variants that likely affect splicing in the *DMD* gene. Among them, 559 (65%) were missense, 131 (15%) were synonymous, 159 (18%) were nonsense, and 14 (2%) were located in untranslated regions (Figure 1-A, 2). Experimental studies are required to determine whether the pathogenicity of these variants is associated with changes at the RNA or protein level.

Interestingly, the majority (98%) of the variants predicted by SpliceAI as potentially affecting splicing were not observed in the general population according to the Genome Aggregation Database (gnomAD). Only 161 predicted SNVs were found in the population data, and among them, 145 had an allelic frequency <0.0003 (the threshold for X-linked diseases) and were not observed in the hemizygous state. These data support that the predicted variants can be considered pathogenic if found in a patient with dystrophinopathy and other necessary ACMG criteria are met. An additional 16 variants had a high allelic frequency; most of these variants were located in deep intronic regions and had relatively low (<0.4) SpliceAI delta scores. Thus, we assume that these are false positive predictions, which is consistent with the reported SpliceAI prediction accuracy.

In summary, using the SpliceAI tool, we predicted a total of 7,085 intronic and 863 exonic variants in the *DMD* gene that could potentially affect pre-mRNA splicing and contribute to dystrophinopathy. Out of these, only 450 variants have been previously described in patients with BMD/DMD and reported in the literature. Thus, we have identified approximately 7,000 variants that likely affect splicing but have not yet been found in individuals with dystrophinopathies.

### Previously reported DMD splicing variants

To determine the presence of SpliceAI-predicted splicing variants in patients with dystrophinopathies, we compared them with disease-associated *DMD* variants previously reported in the scientific literature. At the time of our study, more than half of the reported *DMD* variants were large deletions, insertions, and complex rearrangements. Additionally, ∼19% of the variants were small insertions, deletions, and indels, most of which resulted in frameshifts (Figure 1-B). In the present study, we focused specifically on single nucleotide variants (SNVs), which constituted 27% of all reported *DMD* variants. This included 373 intronic and 956 exonic SNVs. Notably, only about 5% of these variants have previously been functionally characterized.

Of the 373 previously reported intronic SNVs, 353 were assessed by SpliceAI as likely splice-affecting (Figure 2). As expected, the majority of these variants (266 SNVs, 75%) were located at canonical AG-GT terminal dinucleotides of the intron . Only 61 variants (17%) were found at other nucleotides within the acceptor and donor splicing sites. The remaining 26 intronic variants were located outside the splicing sites: one was a +12 variant, while the 25 others were located in deep intronic regions (Figure 1-C). These variants are predicted to result in the creation of donor (19 SNVs) or acceptor (7 SNVs) splicing sites with a SpliceAI delta score range 0.32–0.98. Notably, all 25 deep intronic variants were detected through RNA analysis with subsequent sequence of the target intronic region of the patient’s DNA. Thus, for these variants, it has been experimentally demonstrated that they affect splicing and lead to the creation of pseudoexons. Some of these variants contribute to severe muscular dystrophy phenotypes, while others lead to milder phenotypes (24–36).

**Figure 2.**
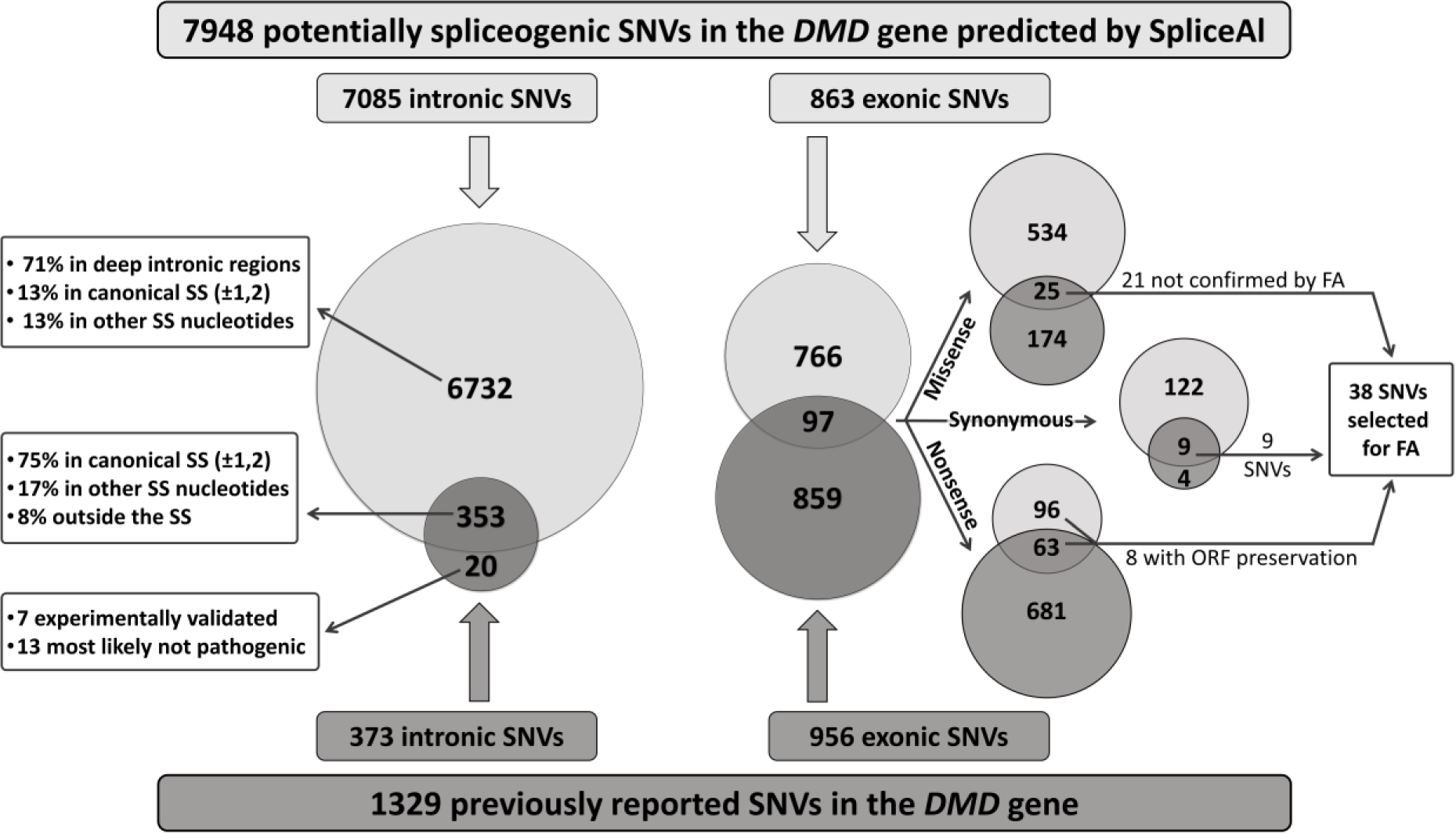
*In silico* research overview. In the present study we analyzed the intersection between all potential splicing-affected *DMD* single nucleotide variants (SNVs) predicted by SpliceAI and all *DMD* SNVs previously reported in the literature. Left: the overlap between the predicted and previously reported intronic SNVs in the *DMD* gene, along with an analysis of their positioning relative to splicing sites (represented in boxes). Right: the overlap between predicted and previously reported exonic SNVs in the *DMD* gene is illustrated for all variants (depicted as large circles) and categorized by variant type (missense, synonymous, nonsense) (depicted as small circles); from each group of overlapping exonic variants, those meeting specific criteria were selected for subsequent experimental functional study. *Abbreviations: SS, Splicing site; FA, Functional analysis; ORF, Open reading frame*.

For the 20 of 373 previously reported intronic SNVs, SpliceAI did not predict a significant effect on splicing (Figure 2). Previously reported RNA analysis revealed that six of these variants (all located in deep intronic regions) disrupted normal splicing patterns and created pseudoexons (25, 30, 37–40), while one resulted in exon skipping due to branch point disruption (41). Interestingly, for the four experimentally validated variants, SpliceAI predicted a corresponding splicing effect, but with a lower delta score (DS = 0.14–0.2), and only two variants had DS of 0 (c.3603+2053G>C, c.9362-1215A>G).

The remaining 13 intronic SNVs not predicted by SpliceAI to affect splicing are likely not associated with the development of dystrophinopathy. Among these, two variants were described *in cis* with another predicted splicing variants and are not pathogenic by themselves (36, 42). Six variants had an allelic frequency >0.0003 and/or were observed in the hemizygous state in the gnomAD database. The remaining five variants were identified during the sequencing of large cohorts of patient samples and were classified as “splicing” solely based on their localization or certain bioinformatic analyses. All of these variants were located in splicing sites (five in the acceptor and one in the donor), but none of thebioinformatic tools applied (SpliceAI, MMSplice, and SPiP) allowed us to determine whether these variants could actually affect splicing.. Hence, approximately 3% of the published intronic variants might have been incorrectly annotated as disease-causing or likely disease-causing.

Out of the 956 previously reported exonic *DMD* SNVs, 97 variants (9 synonymous, 25 missense, and 63 nonsense) were assessed by SpliceAI as likely splice-affecting (Figure 2). Most of them (70%) were located outside the splice sites consensus sequences at exon/intron boundaries (Figure 1-C). The effect of such variants on splicing requires experimental confirmation, but functional studies have been lacking for the majority of potentially spliceogenic SNVs. Therefore about 10% of exonic SNVs in the *DMD* gene may have been inaccurately annotated as affecting protein function, whereas their pathogenicity may actually be due to splicing changes. Understanding the exact molecular mechanism is crucial to explain and predict disease progression and select appropriate therapy. Experimental studies are necessary to identify the precise molecular mechanism underlying the pathogenicity of these SNVs.

Initially, our deep *in silico* mutagenesis focused exclusively on single nucleotide variants (SNVs), excluding small insertions, deletions, and indels due to the vast diversity of such variants. However, these types of variants accounted for 19% of all reported *DMD* alterations (Figure 1-B), and some of them may also impact splicing. To estimate the number of such small rearrangements that might affect splicing, we performed a SpliceAI analysis on all reported small insertions, deletions, and indels. We found that approximately 9% of these small rearrangements in the *DMD* gene had the potential to affect splicing (SpliceAI DS > 0.2). It is important to pay attention to such variants since splicing alterations can lead to mRNA structure changes that differ significantly from the original DNA rearrangement.

### Selection of the variants for the functional study

Out of the 13 published *DMD* synonymous variants, nine were predicted by SpliceAI to be likely splice-altering with a high delta score (DS > 0.6). Although three of these synonymous SNVs had been functionally characterized before and their effect on splicing established, we decided to include all nine variants in our study due to the intriguing mechanism of pathogenicity associated with synonymous variants (Figure 2). The remaining four synonymous variants were not predicted by SpliceAI or other bioinformatics tools to affect splicing. Also, the published reports lacked convincing evidence of their pathogenicity, leading us to exclude these variants from our study (37, 43–45).

Among the 199 published *DMD* missense variants, SpliceAI predicted 25 variants to potentially affect splicing. However, functional confirmation of their splicing effect had been performed only for four variants (46–49). Therefore, we included the remaining 21 missense variants in our study (Figure 2).

Regarding the 744 published nonsense variants, 63 variants overlapped with likely spliceogenic variants predicted by SpliceAI. However, among all types of exonic variants, the pathogenicity of nonsense variants is the most obvious. Therefore, we further analysed all predicted SNVs to find such variants that can lead to the skipping of the whole or part of the exon containing the stop codon formed. Moreover, if the skipped exon is a multiple of three, the reading frame will not shift. Thus, the elimination of the stop codon formed as a result of the nonsense mutation from the mature RNA will lead to completely different consequences at the protein level than the nonsense variant. We identified two such nonsense variants that were previously found in real patients with dystrophinopathies, and an additional six hypothetical variants were predicted by bioinformatics tools. Consequently, we selected these eight nonsense variants for our experimental study (Figure 2).

In summary, a total of 38 exonic *DMD* variants (9 synonymous, 21 missense, 8 nonsense) were selected for the experimental study. To assess the impact of these variants on splicing, we employed a minigene assay as described in the following sections.

### Design and testing of the minigene expression system

To evaluate the effect of exonic *DMD* variants on splicing, we performed a minigene-based assay. This involved cloning the exon (or two exons) of interest with the flanking intronic regions (250– 500 bp of each intron) into the intron of the pSpl3-Flu2-TK vector. All SNVs selected for the study (32 previously published and six theoretical) were located within 27 *DMD* exons. We created 25 minigene constructions with most containing a single exon (23 out of 25), while two constructions contained two exons with a full-length intron between them (Figure 3-A).

**Figure 3.**
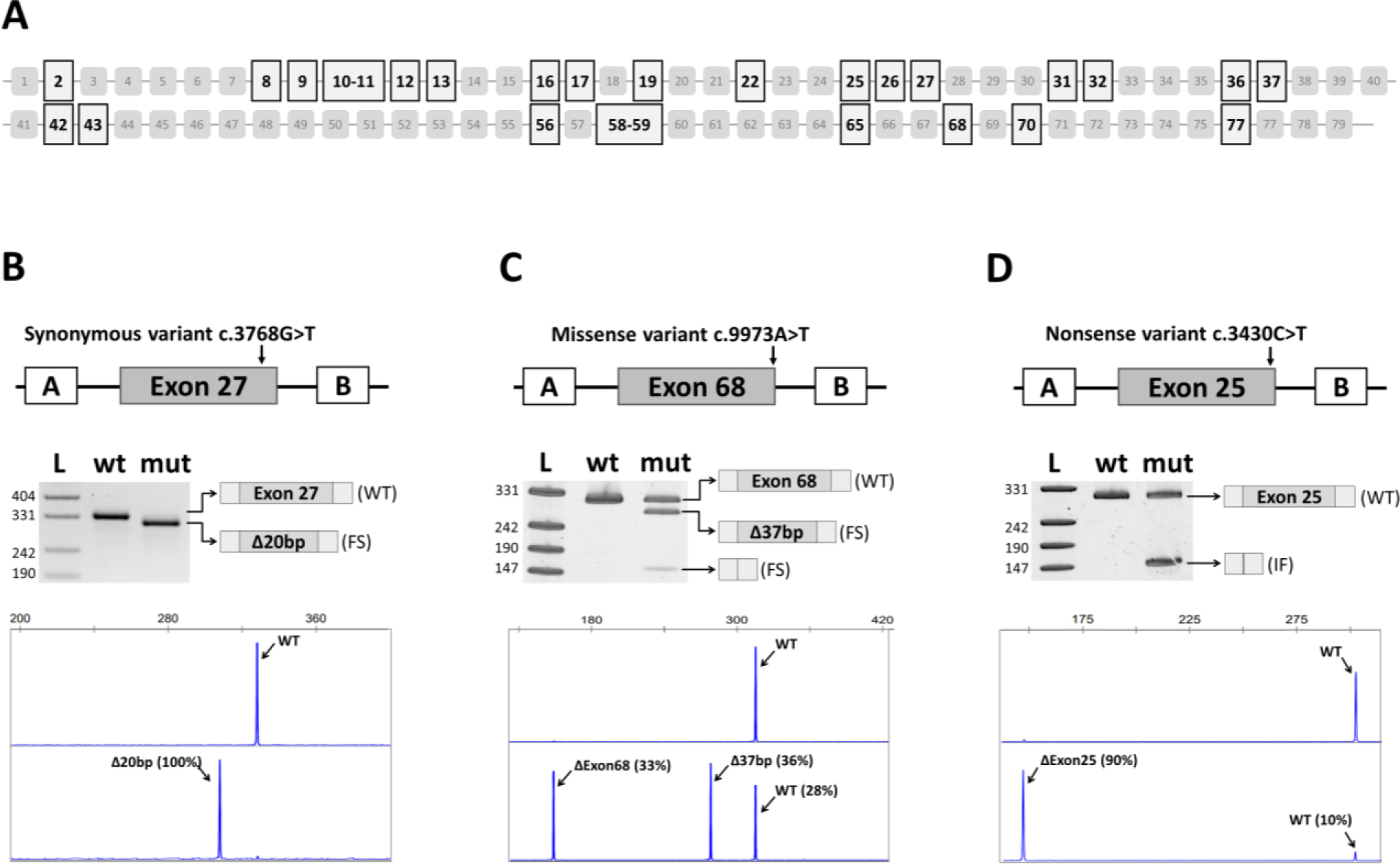
A. Schematic representation of the exonic structure of the *DMD* gene. Exons included in the developed minigene expression system are highlighted with a black frame. B. An example of the functional analysis of synonymous SNVs. The c.3603G>A variant leads to the out-of-frame 20-nt shortening of exon 27 resulting in a PTC. C. An example of the functional analysis of missense SNVs. The c.9973A>T variant leads to the multiple splicing abnormalities. D. An example of the functional analysis of nonsense SNVs. The c.3430C>T variant leads to the in-frame skipping of exon 25 along with residual expression of the full-length transcript. Figures 3B-3D show from top to bottom: scheme of the minigene construction containing the variant studied; electrophoresis of RT-PCR products obtained from minigene transfected HEK293 cells and schematic of the observed splicing changes; fragment analysis of RT-PCR products for a variant-containing plasmid*. Abbreviations: wt, wild type; mut, mutant; FS, frameshift; IF, in-fame;* Δ, *deletion*.

To confirm the correct inclusion of cloned exons we transfected obtained plasmids into HEK293T cells, then performed total RNA extraction and RT-PCR analysis followed by Sanger sequencing. Previous deep sequencing of *DMD* cDNA indicated that the majority of *DMD* exons are constitutively spliced in skeletal muscle under physiological conditions. Although a few non-canonical splicing events were observed, none affected the exons under study, except for the skipping of exon 9 in 0.63% of the transcripts. Our analysis revealed that 20 out of 25 wild-type constructions expressed the expected full-length mRNA. However, five minigenes expressing exons 17, 25, 37, 43 and 70, exhibited incorrect splicing. In two cases we observed partial skipping of the target exons, while three cases displayed aberrantly spliced transcripts not corresponding to any described *DMD* non-canonical splicing events (44).

To ensure correct exon inclusion in these five minigene constructions, we designed seven new minigene plasmids by modifying the length of the flanking intronic regions. Introns are known to contain splicing regulatory motifs that can influence exon inclusion. By this approach we achieved the correct exon inclusion in four of the five cases (exons 17, 25, 37 and 43). . But in the fifth case, along with the full-length transcript, we still observed an aberrant shortening of exon 70. Detailed analysis revealed that this abnormal splicing is due to a weak cryptic splice site located 101 nt downstream of the start of exon 70. Since the wild-type isoform remained one of the primary splicing outcomes, we decided to use this variant of the minigene construction for our further experimental analysis.

In conclusion, we created 25 minigene constructions containing 27 *DMD* exons and tested them in HEK293 eukaryotic cells (Figure S1). Subsequently, the selected 38 SNVs were introduced into the wild-type minigene plasmids using site-directed mutagenesis, and we conducted experimental studies to examine the effect of these variants on pre-mRNA splicing.

### Functional analysis of synonymous variants

Among the nine selected synonymous SNVs, five variants were located in the last nucleotide of the exon, which, according to bioinformatics predictions, should lead to the loss of the donor splice site. Our functional analysis confirmed that all five variants indeed impair splicing. As a result, we observed the appearance of one (NM_004006.2:c.4518G>A, c.3432G>A) or several aberrant transcripts (c.6117G>A, c.3603G>A, c.1602G>A) due to complete or partial exon skipping and/or intron retention caused by the activation of cryptic splicing sites in the intron (Figure 3-B, Figure S2).

The remaining four synonymous variants are located upstream of the last nucleotide of the exon (c.4299G>T, c.3768G>T, c.1329C>T, c.1098A>T). Bioinformatics predictions suggested that these variants should lead to the formation of a new donor splice site. Functional analysis confirmed that the use of new splice sites in all 4 cases leads to exon truncation followed by a frameshift. Three of the synonymous variants under study (c.6117G>A, c.3432G>A, c.1602G>A) had previously been investigated by RT-PCR analysis of the affected individual’s tissues (24, 47, 50). Our minigene-based analysis of these variants confirmed qualitative splicing impairment, consistent with the previously reported results. This finding further validates the reliability and concordance of the minigene assay with RT-PCR. However, in the case of variant c.1602G>A, previous RT-PCR data showed exon 13 skipping in the patient’s muscle biopsy (50). In our minigene analysis, in addition to exon skipping, we also observed residual expression of the wild-type isoform. This discrepancy may be attributed to limitations of the minigene system as well as tissue-specific differences.

In summary, our analysis demonstrated that all nine synonymous variants actually affect pre-mRNA splicing (Table 1). Six out of the nine variants lead toa frameshift and inclusion of a premature termination codon. Moreover, in three cases (c.3768G>T, c.1329C>T, c.1098A>T), the aberrant transcript was the only isoform observed. Therefore, these SNVs can be classified as loss-of-function variants and should be considered likely pathogenic according to the ACMG criteria. The remaining three variants, along with the isoform containing a premature stop codon, retained residual expression of the wild-type and/or in-frame isoforms, suggesting milder molecular consequences.

**Table 1.** Splicing outcomes of *DMD* synonymous SNVs.

| SNV* | PMID | Phenotype | Exon | SpliceAI | Observed splicing events (%) |  |  |
| --- | --- | --- | --- | --- | --- | --- | --- |
|  |  |  |  |  | in frame | out of frame | WT |
| c.6117G>A | 20485447 | BMD | 42 | DL: 0.72;<br>DG: 0.39 | Exon skipping (12%) | 35-nt intron retention (81%) | 7% |
| c.4518G>A | 19937601 | IMD | 32 | DL: 0.62;<br>DG: 0.54 | Exon skipping (100%) | - | - |
| c.4299G>T | 32194622 | BMD | 31 | DG: 0.48;<br>DL: 0.59 | Exon skipping (4.5%) | 47-nt deletion (87%) | 8,5% |
| c.3768G>T | 32194622 | BMD | 27 | DG: 0.99;<br>DL: 0.74 | - | 20-nt deletion (100%) | - |
| c.3603G>A | 19959795 | DMD | 26 | DL: 0.89 | Exon skipping (4%) | Intron retention (71%) | 23% |
| c.3432G>A | 19783145 | BMD | 25 | DL: 0.98 | Exon skipping (100%) | - | - |
| c.1602G>A | 31443951;<br>31706698 | BMD | 13 | DL: 0.91 | Exon skipping;<br>90-nt deletion (15%) | - | 85% |
| c.1329C>T | 32559196 | BMD | 11 | DG: 0.98;<br>DL: 0.61 | - | 4-nt deletion (100%) | - |
| c.1098A>T | 11524473 | DMD | 10 | DG: 0.98 | - | 53-nt deletion (100%) | - |

\*Variants were named following Human Genome Variation Society (HGVS) guidelines with reference NM\_004006.2. Abbreviations: WT, wild-type; BMD, Becker muscular dystrophy; IMD, Intermediate muscular dystrophy; DMD, Duchenne muscular dystrophy; DG, Donor gain; DL, Donor loss; nt, nucleotide.

### Functional analysis of missense variants

Among the 21 missense variants selected, three variants according to bioinformatics predictions should lead to the formation of a new acceptor splice site. Our functional analysis revealed that two of these three variants indeed impaired normal splicing and caused shortening of corresponding exon (Figure S3). The variant c.2827C>T showed no effect on splicing and had a relatively low SpliceAI delta score (DS = 0.2687). It should be noted that this variant was found in a patient with Stargardt macular dystrophy and not associated with dystrophinopathies, as reported by Zaneveld et al. (51).

Twelve out of the 21 selected missense variants were located in the last or second-to-last nucleotide of the exon, potentially leading to the loss of the donor site. Our functional analysis demonstrated that all twelve variants caused splicing abnormalities, including complete or partial exon skipping, intron retention, and more complex splicing aberrations such as activation of cryptic sites or the appearance of multiple mis-spliced isoforms (Figure S3).

According to SpliceAI predictions, seven out of the 21 selected missense variants located deeper in exons were expected to lead to the formation of a new donor splice site. Our functional analysis confirmed that six out of the seven variants indeed affected splicing and resulted in the appearance of one or several aberrant transcripts. The variant c.1934A>G showed no effect on splicing and had a relatively low SpliceAI delta score (DS = 0.3449).

In conclusion, our functional analysis revealed that 19 out of the 21 tested missense variants in fact affect splicing (Table 2). Among these variants, ten led to the production of one or several aberrantly spliced transcripts. Remaining nine variants alongside the aberrant transcripts produced a various amount of the wild-type isoform (Figure 3-C, Figure S3).

**Table 2.** Splicing outcomes of *DMD* missense SNVs.

| SNV | PMID | Phenotype | Exon | SpliceAI | Observed splicing events (%) |  |  |
| --- | --- | --- | --- | --- | --- | --- | --- |
|  |  |  |  |  | in frame | out of frame | WT |
| c.10149A>C | 19937601;<br>25220406 | B/DMD | 70 | AG: 0.84 | - | 65-nt deletion (85%) | 15% |
| c.9973A>T | 32169422 | DP | 68 | DL: 0.42;<br>DG: 0.26 | - | Exon skipping;<br>37-nt deletion;<br>Intron retention (72%) | 28% |
| c.9560A>G | 17259292 | DMD | 65 | DG: 0.99;<br>DL: 0.70 | - | 4-nt deletion (100%) | - |
| c.8937G>C | 27593222 | MD | 59 | DL: 0.79 | 75-nt deletion (96%) | - | 4% |
| c.8668G>A | 22776072;<br>25220406 | BMD | 58 | DL: 0.41 | - | Exon skipping (80%) | 20% |
| c.8390G>A | 22776072 | BMD | 56 | DL: 0.77 | - | Intron retention (17%) | 83% |
| c.8390G>C | 11524473 | DMD | 56 | DL: 0.97 | - | Exon skipping;<br>Intron retention (100%) | - |
| c.6290G>T | 30907348 | DMD | 43 | DG: 0.61 | - | 4-nt intron retention (50%) | 50% |
| c.6117G>C | 19937601;<br>25525159 | DMD | 42 | DL: 0.87;<br>DG: 0.45 | Exon skipping (6%) | Intron retention (94%) | - |
| c.5154G>T | 32528171 | LGMD | 36 | DL: 0.98;<br>DG: 0.31 | Exon skipping (54%) | Intron retention (46%) | - |
| c.3432G>T | 23756440;<br>29604111 | BMD | 25 | DL: 0.98 | Exon skipping (100%) | - | - |
| c.2827C>T | 25474345 | STGD | 22 | AG: 0.27 | - | - | 100% |
| c.2949G>T | 28859693;<br>29973226 | DMD | 22 | DL: 0.87 | - | Exon skipping (90%) | 10% |
| c.2378A>G | 22776072;<br>25220406 | BMD | 19 | DG: 0.98;<br>DL: 0.71 | - | 7-nt deletion (100%) | - |
| c.2380G>A | 29365344 | DMD | 19 | DL: 0.89;<br>DG: 0.56 | - | Exon skipping;<br>7-nt deletion (97%) | 3% |
| c.1934A>G | 7981690 | DMD | 16 | DG: 0.35 | - | - | 100% |
| c.1351G>T | 22776072;<br>32597815 | BMD | 12 | AG: 0.87 | - | 28-nt deletion (100%) | - |
| c.1310C>G | 19937601;<br>32559196 | DMD | 11 | DG: 0.96 | - | 26-nt deletion (100%) | - |
| c.1149G>T | 27122458 | B/DMD | 10 | DL: 0.72 | - | 50/274-nt intron retention (62%) | 38% |
| c.831G>T | 30833962 | DMD | 8 | DL: 0.99 | 105-nt deletion (53%) | Exon skipping;<br>5/61-nt intron retention (47%) | - |
| c.77A>G | 30833962;<br>31081998 | DMD | 2 | DG: 0.92 | 21-nt deletion (100%) | - | - |

Abbreviations: WT, wild-type; B/DMD, Becker/Duchenne muscular dystrophy (Unknown phenotype); MD, Muscular dystrophy; BMD, Becker muscular dystrophy; DP, Dystrophinopathy; LGMD, Limb girdle muscular dystrophy; DMD, Duchenne muscular dystrophy; STGD, Stargardt macular dystrophy; AG, Acceptor gain; DG, Donor gain; DL, Donor loss; nt, nucleotide.

### Functional analysis of nonsense variants

Among the eight selected nonsense variants, only one SNV did not showed any effect on splicing in our functional analysis (Table 3). Seven out of the eight nonsense variants led to various splicing aberrations, including loss of the donor splice site and the formation of a new and/or activation of a cryptic donor splice site. These events resulted in partial or complete exon skipping and/or intron retention. However, in all of these cases, we observed variable expression of the full-length mRNA isoform, indicating that these variants can lead to the synthesis of truncated proteins in different amounts.

**Table 3.** Splicing outcomes of *DMD* nonsense SNVs.

| SNV | PMID | Phenotype | Ex on | SpliceAI | Observed splicing events (%) |  |  |
| --- | --- | --- | --- | --- | --- | --- | --- |
|  |  |  |  |  | in frame | out of frame | WT |
| c.11014G>T | - | Hypothetic | 77 | DL: 0.46 | Exon skipping (82%) | - | 18% |
| c.8668G>T | - | Hypothetic | 58 | DL: 0.6 | - | Exon skipping (90%) | 10% |
| c.5323A>T | - | Hypothetic | 37 | DL: 0.64 | Exon skipping (100%) | - | - |
| c.3438T>G | - | Hypothetic | 26 | AG: 0.78 | - | - | 100% |
| c.3430C>T | 19959795 | DMD | 25 | DL: 0.55 | Exon skipping (90%) | - | 10% |
| c.2380G>T | 11524473 | DMD | 19 | DG: 0.25;<br>DL: 0.8 | - | Exon skipping;<br>7-nt deletion (36%) | 64% |
| c.2164A>T | - | Hypothetic | 17 | DG: 0.67 | 6-nt deletion (9%) | Intron retention (7%) | 84% |
| c.958C>T | - | Hypothetic | 9 | DL: 0.56 | Exon skipping (56%) | - | 34% |

*Abbreviations: WT, wild-type; DMD, Duchenne muscular dystrophy; DG, Donor gain; DL, Donor loss; nt, nucleotide.*

In six out of the seven cases, exon skipping was the predominant mutant isoform due to the loss of the donor splice site. Notably, in four cases (c.11014G>T, c.5323A>T, c.3430C>T, c.958C>T), the skipped exon was a multiple of three, preserving the reading frame. Consequently, instead of a premature stop codon, these variants led to an in-frame deletion, significantly altering their molecular consequences and the underlying mechanism of pathogenicity (Figure 3-D, Figure S4).

For two variants (c.3438T>G and c.2164A>T), we expected the SNV to lead to the creation of a new splice site, resulting in localization of the premature stop codon within an intronic sequence recognized by the spliceosome. Our functional analysis demonstrated that variant c.2164A>T caused an in-frame deletion of the exon region containing the nonsense variant-induced stop codon. However, the proportion of this isoform was only 6% of the total number of transcripts. Variant c.3438T>G, despite its high SpliceAI delta score (DS = 0.78), did not alter splicing and can be considered a true nonsense variant.

### Analysis of molecular consequences of splicing SNVs to explain observed DMD and BMD phenotypes

Our investigation into the molecular consequences of *DMD* variants on splicing has provided valuable insights into their relationship with patient phenotypes. To establish this connection, we compiled available information on patients carrying the variants under study. However, it is important to note that the amount of available information varied significantly. Some publications lacking additional details beyond the diagnosis (DMD or BMD) because they screened a large cohort of patients using next-generation sequencing methods. (Table S3).

Next, we analyzed all splicing events identified during the functional analysis of exonic variants in the *DMD* gene, specifically focusing on synonymous and missense variants with splicing impact. We categorized the isoforms detected in each case into three groups: full-length wild-type isoform, in-frame aberrant isoform, and out-of-frame aberrant isoform. Subsequently, we calculated the proportions of each isoform type and compared their relative abundance between two patient groups: Group 1 consisting of individuals with the DMD phenotype, and Group 2 comprising patients with milder phenotypes such as BMD, intermediate forms (IMD, MD), or other related diagnoses (Dystrophinopathy, Muscular dystrophy, limb-girdle,). For two variants with the phenotype described as B/DMD, no clarifying data could be found, so they were excluded from the analysis.

Our findings revealed a significant increase in the level of out-of-frame aberrant isoforms in patients exhibiting the DMD phenotype (Group 1). Conversely, patients with milder phenotypes (Group 2) showed significantly higher levels of wild-type and in-frame isoforms. This trend suggests a correlation between the molecular genetic cause of the disease and the observed patient phenotype (Figure 4-A).

**Figure 4.**
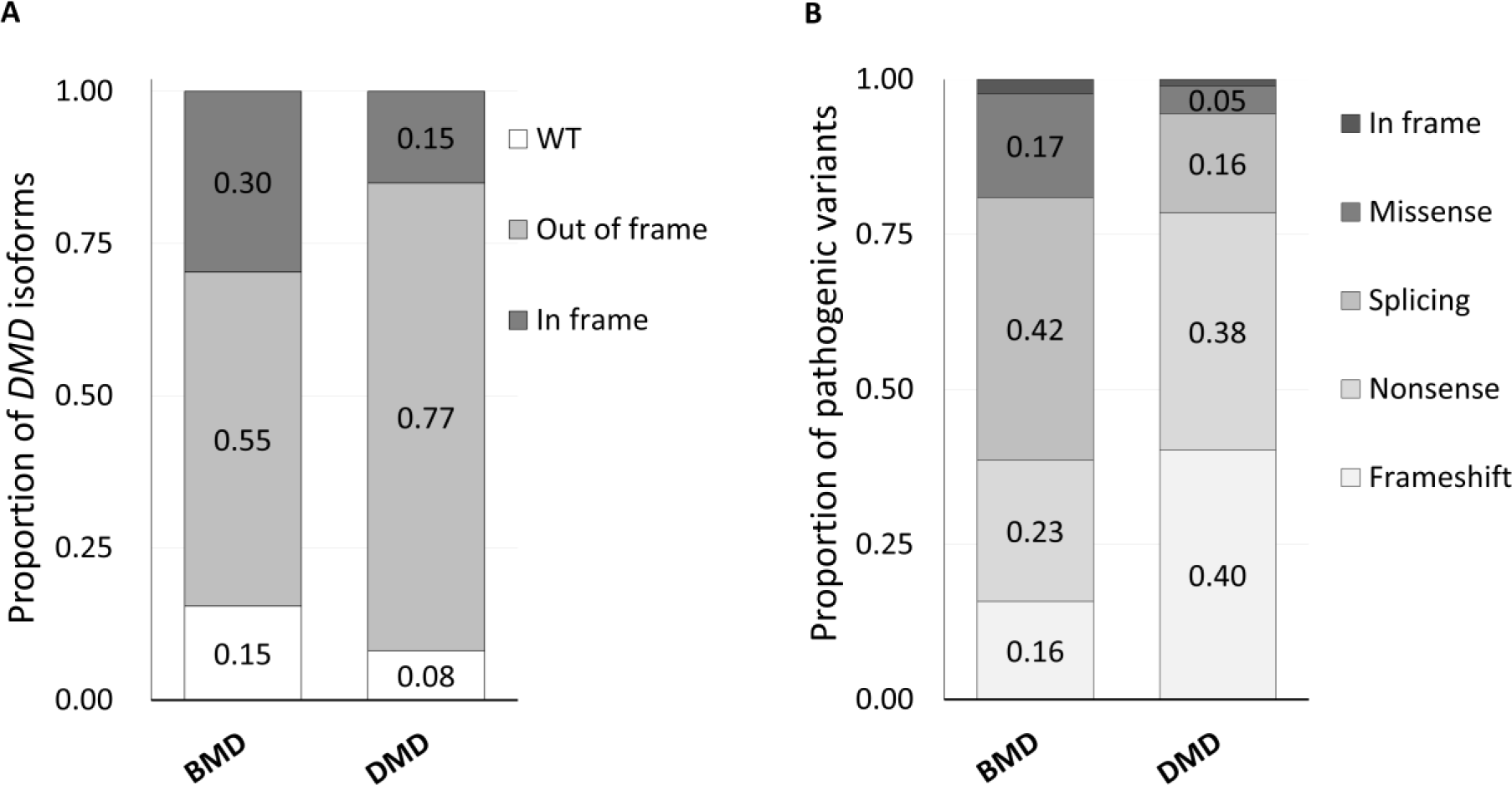
A. Relative abundance of the wild-type, in-frame-, and out-of-frame isoforms derived from experimental analysis of previously reported exonic *DMD* variants using our minigene system. Variations in the isoform proportions were detected between DMD (Duchenne muscular dystrophy) and BMD (Becker muscular dystrophy) patient-identified variants. B. Relative abundance of the different types of pathogenic variants in the *DMD* gene found in patients with muscular dystrophies, described in the literature.

To further enhance our understanding of the association between different groups of pathogenic variants in the *DMD* gene and phenotypes, as well as to compare our results with existing data, we performed an analysis of pathogenic variants reported in the literature for BMD (n = 215) and DMD (n = 1442) patients. Our analysis revealed that frameshift and nonsense variants accounted for over 75% of all pathogenic variants identified in DMD patients. In contrast, the proportion of such variants in BMD patients was less than 40%, with a higher prevalence of splicing, missense, and in-frame variants (Figure 4-B).

Thus, by integrating our own findings with the broader data, we have extended our insights into how the molecular mechanisms of pathogenicity of splicing variants are linked to the clinical features observed in patients with DMD and BMD.

## Discussion

Pathogenic variants affecting pre-mRNA splicing constitute a distinct class of disease-causing genetic variants. Estimates suggest that splicing variants are associated with 30% to 50% of hereditary diseases (52, 53). These variants can directly cause the disease, influence disease susceptibility, or modulate disease severity by affecting canonical splice sites, creating new splice sites, or altering splicing regulation (54).

Despite the considerable number of splicing variants reported in the DMD gene, large-scale studies on splicing in this gene are lacking. There are a number of studies reporting the functional analysis of several distinct splicing variants, but they do not provide a complete picture of the place of splicing variants in the structure of all pathogenic *DMD* variants. Most studies have focused on variants located at consensus positions of splice sites and have rarely explored exonic splicing variants (10, 68, 69). Moreover, when a newly identified variant is reported to affect splicing, it is often based solely on bioinformatics predictions without experimental confirmation (45, 70) In this study, we attempted to characterize the full spectrum of splicing variants in the *DMD* gene. All SNVs previously reported in the literature, as well as all possible single-nucleotide substitutions across the entire *DMD* gene sequence were analyzed for their effect on splicing. This analysis provides the most comprehensive view of the number, location, and distribution of the splicing variants in the DMD gene.

One of the expected findings of this analysis is the significant underrepresentation of splicing variants among both intronic and exonic SNVs identified in patients with dystrophinopathies. At the same time, some patients do not have an identified causative *DMD* variant. We found that the majority of intronic splicing variants reported to date in the *DMD* gene are located in the canonical AG-GT dinucleotides of the splicing sites. However, unbiased SpliceAI evaluation predicted a nearly equal number of variants potentially affecting splicing at terminal dinucleotides and other positions within the splicing sites, even when using a stringent SpliceAI DS > 0.5 threshold. Moreover, SpliceAI estimates that more than 60% of all possible likely splice-affecting variants are located deep in introns of the *DMD* gene. Although only 25 deep intronic *DMD* splicing variants have been previously described in the literature. It is noteworthy that two recent independent studies of causative variants in genetically undiagnosed patients with dystrophin deficiency in muscle showed that the majority of these patients carry variants located deep in introns (71, 72).

Our in-silico analysis of exonic SNVs also revealed the significant discrepancies between predicted and previously reported variants. Although the SpliceAI tool has some inherent limitations and we acknowledge the potential for bias arising from its training data, we consider its predictions to be highly credible. Therefore, we suggest that routine DNA diagnostics primarily focus on canonical terminal AG-GT dinucleotide variants, while other variants are often overlooked.

The challenges in detecting splicing variants in the *DMD* gene arise from the current standards for genetic diagnosis of dystrophinopathies. Since the majority of patients exhibit exon deletions or duplications, multiplex ligation-dependent probe amplification (MLPA) is the most cost- and labor-efficient method for initial molecular diagnosis (73, 74). Small mutations require Sanger sequencing or next-generation sequencing (NGS), such as targeted sequencing of the *DMD* gene or whole exome sequencing (WES). However, these methods may not reveal certain splicing alterations, particularly deep intronic variants since target panels and WES do not cover intronic sequences. Additionally, the impact of most synonymous and missense variants on splicing mechanisms often remains challenging to determine, as the main focus is typically on protein structure and function changes, with potential effects on splicing being underestimated (75).

Experimental verification is crucial for definitively diagnosing splicing variants. In recent years, reverse transcription-polymerase chain reaction (RT-PCR) and experimental confirmation using minigenes have been the primary methods for testing splicing variants. However, RT-PCR analysis is limited by the availability of patient tissue samples, and mRNA degradation via the nonsense-mediated decay (NMD) mechanism may hinder detection of aberrant isoforms. Minigene assays, on the other hand, allow the creation of variants *in vitro* using site-directed mutagenesis and do not rely on patient samples. However, they may lack the broader genomic context and may not fully reflect the splicing pattern in patients, necessitating careful selection of the system for study.

In this study, we performed the minigene-based analysis of 38 exonic single nucleotide variants (SNVs) in the *DMD* gene to assess their effect on splicing. Our analysis allowed to reclassify synonymous and missense variants that affect splicing from the category of variants of uncertain significance (VUS) to likely pathogenic (PS3, PM2). It is noteworthy that out of the 38 variants selected for functional analysis, only 3 did not exhibit any effect on splicing. Specifically, the missense variants c.2827C>T and c.1934A>G had relatively low delta scores (DS = 0.2687 and 0.3449, respectively), which might have caused them to be overlooked if the threshold had been set higher. The nonsense variant c.3438T>G was bioinformatically predicted to create an acceptor site, but our analysis revealed that the wild-type site was used instead. These data do not contradict the expected accuracy of SpliсeAI (23, 76). Moreover, SpliceAI predicted the creation of a new acceptor site but not the disruption of a wild-type site. According to MAXENTSCAN, the strength of the new site is significantly lower than the strength of the wild-type site (MAXENT: 5.20 and -2.74, respectively), therefore, the wild-type site can be used, as shown by our analysis. Additionally, we evaluated the c.3438T>G variant using the MMSplice and SPiP tools, which also did not indicate its impact on splicing.

Understanding the molecular mechanism of pathogenicity for an SNV found in a patient is crucial for explaining its phenotype. The “reading-frame rule” can explain approximately 90% of phenotypes associated with large rearrangements in the *DMD* gene (79). However, around 10% of DMD/BMD cases caused by point mutations do not follow this rule. While some exceptions are described in the literature, the explanation for many of these events remains unknown (12, 80, 81). Splicing variants can also lead to exon skipping, intron retention, and other events that affect the reading frame. Splicing aberrations that maintain the reading frame may result in a milder phenotype compared to frameshift variants that lead to the loss of protein function. Moreover, in-frame splicing changes can still be loss-of-function variants if they affect important structural or catalytic domains. These factors often complicate the assessment of the pathogenicity of splicing variants.

To evaluate the correspondence of our results with the reading-frame rule, we examined published data on the phenotypes of patients carrying the variants selected for our functional study. Our analysis showed that the proportion of out-of-frame isoforms in DMD patients is ∼80%. On the other hand, nearly half (∼45%) of the total number of isoforms in patients with BMD are in-frame and wild-type. These findings do not contradict the reading frame rule and align with previous studies (9, 77, 78).

The mechanism of pathogenicity of the synonymous variants is often not obvious and such variants can be misclassified as non-pathogenic. However, several cases have been described where a synonymous variant resulted in complete or partial exon skipping, leading to various diseases (79, 80). In our study, all nine tested synonymous variants were found to affect pre-RNA splicing, confirming the utility of functional analysis for molecular diagnosis in cases where no other pathogenic variants in the *DMD* gene are identified.

The pathogenicity of missense variants is usually associated with amino acid substitutions that render the protein nonfunctional. However, depending on the properties of the amino acid or its location, missense variants may not affect protein function. Although relatively few pathogenic missense variants in the *DMD* gene have been described to date (50, 81), they are predominantly localized in two key functional domains of dystrophin. The most common pathogenic missense variants in the N-terminal actin-binding domain, associated with the milder BMD phenotype, can disrupt actin-binding activity (82), compromise dystrophin’s thermal stability, and increase its propensity for aggregation *in vitro* (81, 83). Variants in the ZZ β-dystroglycan binding domain, which typically causes the more severe DMD phenotype, abolish the binding to β-dystroglycan, which in turn connects to the extracellular matrix (84, 85). Nevertheless, for a variety of the missense variants located throughout the *DMD* gene sequence, assigning pathogenicity is difficult and they often remain variants of uncertain significance (86).

Like synonymous SNVs, the missense variants localized in the splice site can led to splicing abnormalities and disrupt protein function. For example, the missense variant c.1310C>G leads to the formation of a new donor site and deletion of 26 nt of exon 11. The subsequent frameshift and premature stop codon can explain the severe phenotype observed in the affected individual, which is described as DMD (9). On the other hand, the missense variant c.9973A>T results in incomplete aberrant splicing, with some amounts of the full-length transcript (28%), which is consistent with milder phenotypes observed in patients according to the study by Kadiri et al. (77).

Interestingly, in almost half (18 out of 38) of the tested single nucleotide variants (SNVs), we observed residual amounts of the wild-type transcript ranging from 4% to 80%. The severity of the phenotype in dystrophinopathy largely depends on the residual amount of dystrophin. Previous studies have demonstrated that dystrophin levels ranging from 29% to 57% are sufficient to avoid significant signs of myopathy, such as muscle weakness (87). Furthermore, de Feraudy et al. showed that even as low as 5-0.5% of full-length dystrophin was associated with milder clinical signs, including delayed loss of ambulation (88). These findings are supported by studies on a mouse model for Duchenne muscular dystrophy, where dystrophin levels below 4% were shown to improve survival and motor function (89).

It is important to note that the significance of residual full-length transcript expression may not be so precise when viewed in terms of exonic variant types. In the case of a synonymous substitution affected splicing, residual amounts of full-length dystrophin will indeed contribute to the mildering of the phenotype, since the amino acid sequence of the full-length transcript is not changed. If splice-affecting missense variant produce residual full-length transcript than it is difficult to assess how the resulting missense substitution will affect protein function. As mentioned above, the most common missense variants in the *DMD* gene are associated with the milder BMD phenotype so in some cases expression of the full-length transcript could moderate the severe consequences of the aberrant splicing. In the case of the splice-altering nonsense variants, the full-length transcript will contain a premature stop codon and its expression can only aggravate the severity of the phenotype. Thus, the significance of residual expression of the full-length transcript should be carefully assessed on a case-by-case basis. In addition to understanding the genotype-phenotype relationship, precise knowledge of the molecular mechanisms underlying the pathogenicity of *DMD* gene variants is crucial for evaluating and selecting appropriate genetic therapeutic options for each patient. Given the progressive nature of dystrophinopathy and the irreversible muscle loss associated with the condition, initiating eligible gene therapy as early as possible is of utmost importance.

A promising genetic therapy for DMD is stop-codon readthrough, which enables the translation mechanism to bypass premature stop codons caused by nonsense mutations (20). Currently, the drug ataluren (PTC Therapeutics “Translarna”) has been approved in Europe for the treatment of DMD caused by nonsense mutations in ambulatory patients aged 5 years or older (90). However, in cases where the nonsense variant leads to the activation of a new splice site or cryptic splice site, the premature stop codon can be excluded from the transcript, rendering expensive therapy with ataluren potentially ineffective.

Another therapeutic approach based on understanding the molecular mechanism of pathogenicity is antisense oligonucleotide (AON)-mediated exon skipping. This method is currently one of the most promising therapeutic strategies for DMD (22). Exon-skipping therapy aims to modify pre-mRNA splicing, resulting in the skipping of one or more exons and restoration of the reading frame (91). As a result, truncated but partially functional dystrophin protein, as seen in BMD, is produced (92). Several drugs targeting different exons are currently being developed to address various DMD patient subpopulations (93, 94). Accurate molecular genetic diagnosis is necessary to determine the suitability of using expensive targeted drugs.

## Conclusion

In this study, we conducted a comprehensive bioinformatics analysis to assess all possible variants that can affect splicing in the *DMD* gene. We developed a minigene expression system to functionally evaluate the impact of exonic single nucleotide variants (SNVs) on splicing. Our findings revealed that among the *DMD* variants previously identified in patients with dystrophinopathies, 19 missense, nine synonymous, and seven nonsense variants actually disrupt splicing. This novel information regarding the molecular mechanisms underlying the pathogenicity of exonic SNVs provides valuable insights into the clinical characteristics of individuals affected by Duchenne and Becker muscular dystrophy.

## Materials and methods

### Variant nomenclature and accession

Single nucleotide variants (SNVs) were named in accordance with the Human Genome Variation Society (HGVS) recommendations (95). The hg19 assembly and the NG_012232.1 were used as reference genome sequences. The NM_004006.2 transcript corresponding to the full-length muscle isoform Dp427m was used as a reference.

Human Genome Mutation Database (HGMD Professional 2022.1) were used to collect all published SNVs affecting splicing in the *DMD* gene.

### Bioinformatic analysis

The effects of SNVs on pre-mRNA splicing were predicted using the SpliceAI (Illumina, USA) (23), MaxEntScan (MIT, USA) (96), and Human Splicing Finder (Aix Marseille Université, France ) (97) tools. Distinct variants were also evaluated using the MMSplice (98) and SPiP (99) tools. Variants were annotated using the wANNOVAR tool (Wang Genomics Lab, USA) (100).

### Minigene constructions

We used the pSpl3-Flu2-TK vector to create minigene plasmids. This vector was derived from the previously described pSpl3-Flu vector (101) by deleting the cryptic splicing sites within the HIV-tat intron and replacing the strong CMV promoter with the weaker HSV TK promoter. The use of the TK promoter leads to the expression of minigenes at a level closer to the physiological level.

Wild-type minigene plasmids were prepared using target exons and variable flanking intronic sequences of the *DMD* gene amplified from a control genomic DNA using Q5 High-Fidelity DNA Polymerase (NEB, USA). The primers that were used are described in Table S1. The PCR products were cloned into the pSpl3-Flu2-TK-del vector using Gibson Assembly Master Mix (NEB, USA).

Mutant minigene plasmids were obtained by introduction of selected SNVs into wild-type plasmids by the single-primer site-directed mutagenesis method (102) or using the Phusion Site-Directed Mutagenesis Kit (Thermo Scientific, USA) (Table S1). The nucleotide sequence of the plasmids was confirmed by Sanger sequencing.

### Transfection of eukaryotic cells

Human embryonic kidney 293T (HEK293T) cells were cultured in high glucose DMEM with alanyl-glutamine (PanEco, Russia) supplemented with 10% fetal bovine serum (Biosera, France) in 5% CO_2_ at 37°C. Twenty-four hours before transfection, a 24-well plate was treated with poly-L-lysine and 6×10^4^ cells were seeded in each well. Two hours before transfection the growth medium was replaced with DMEM without FBS. Samples containing 750 ng of wild-type or mutant plasmid then were transfected into HEK293T cells by the calcium phosphate transfection method (103). For transfection control at least 2 wells of each plate were transfected with an empty pSpl3-Flu2-TK-del plasmid. The empty vector was also used as an internal control for further evaluation of the transfection efficiency (positive control). To control of cell survival and the absence of contamination untreated cells were used (negative control). The efficiency of transfection was assessed using a flow cytometer and the FloMax software package (Partec, Germany).

### RNA splicing assays

Forty-eight hours after transfection, cells were harvested and total RNA was isolated using the ExtractRNA reagent (Evrogen, Russia) according to the manufacturer’s recommendations. The isolated RNA was treated with DNaseI (Thermo Fisher, USA). cDNA was synthesized using the ImProm-II™ Reverse Transcription System (Promega, USA). The mRNA composition was established by PCR with plasmid-specific primers. Previously well-tried cDNA obtained from cells transfected with an empty pSpl3-Flu2-TK-del plasmid was used as a positive control. The PCR products were analysed by denaturing urea polyacrylamide gel electrophoresis followed by Sanger sequencing (Table S1).

### Fragment analysis

cDNA was amplified as described above using a forward primer carrying a 6-FAM tag. The products were separated on an Applied Biosystems 3500/3500xL Genetic Analyzers capillary electrophoresis system (Thermo Fisher, USA). The relative quantities of the isoforms were assessed from the peak heights of each isoform compared to the total area under the curve and peak height for all isoforms for each sample.

## Declarations

### Ethics approval and consent to participate

Not applicable

### Consent for publication

Not applicable

### Authors’ contributions

KD performed experiments, analysed and interpreted the data and was primary contributor to writing the manuscript; AF designed experiments, performed bioinformatics and statistical analysis and revised the manuscript; MS designed the study, analysed and interpreted the data, and revised the manuscript. All authors read and approved the final manuscript.

## Acknowledgements

We are grateful to Ilia Korvigo for participation in part of the work with «In silico analysis of possible splicing SNVs in the DMD gene». We thank Richard H. Lozier for editorial assistance.

## Availability of data and material

All data generated or analysed during this study are included in this published article [and its supplementary information files].

