## Supplemental Figure S1 for "Revision of splicing variants in the *DMD* gene"

| Exon № | Scheme of the locus | Fragment analysis |
|---|---|---|
| 2 | 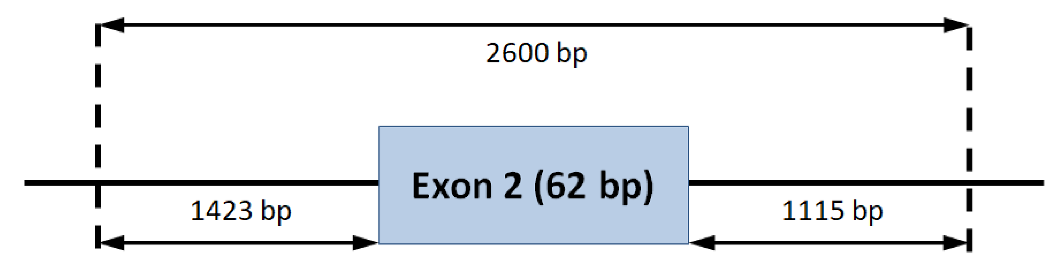 | 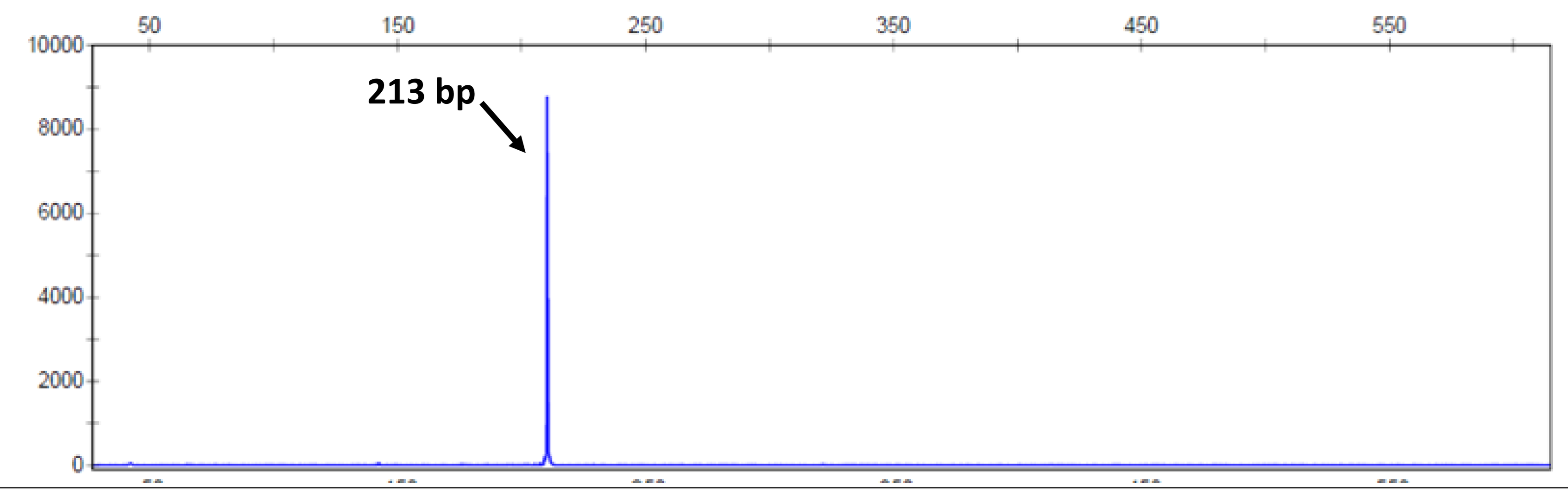 |
| 8 | 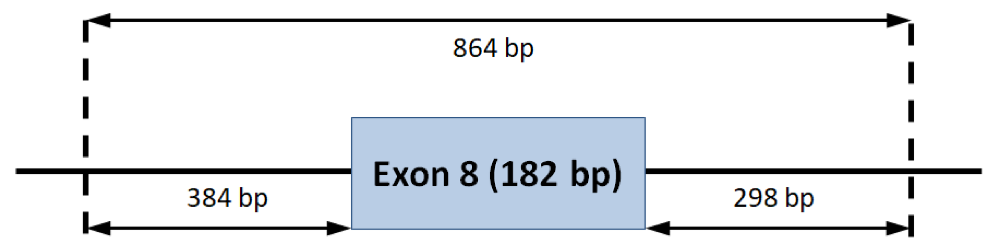 | 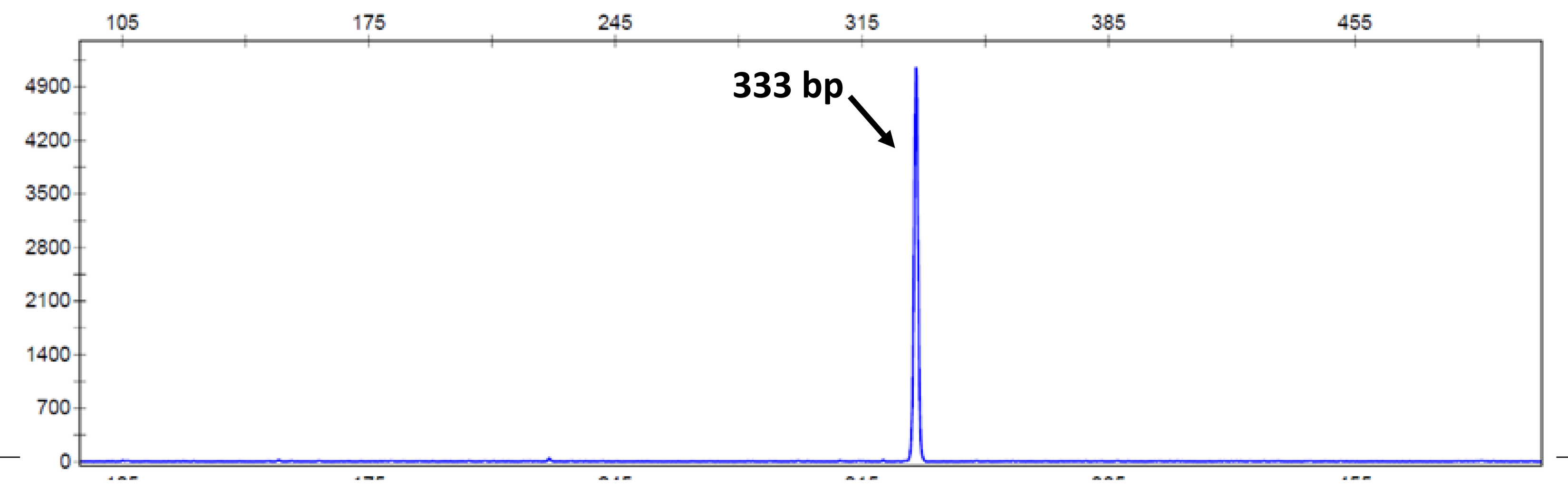 |
| 9 | 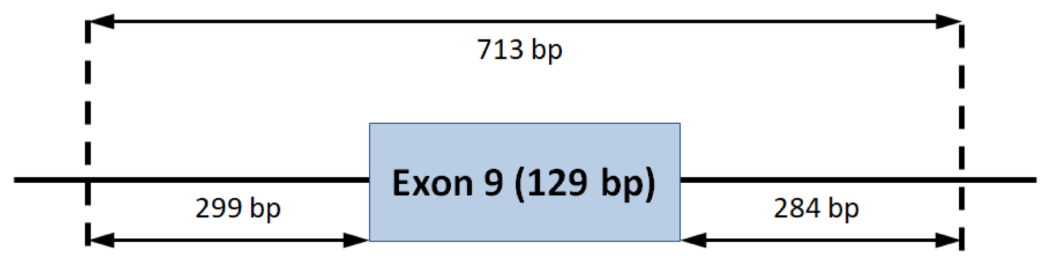 | 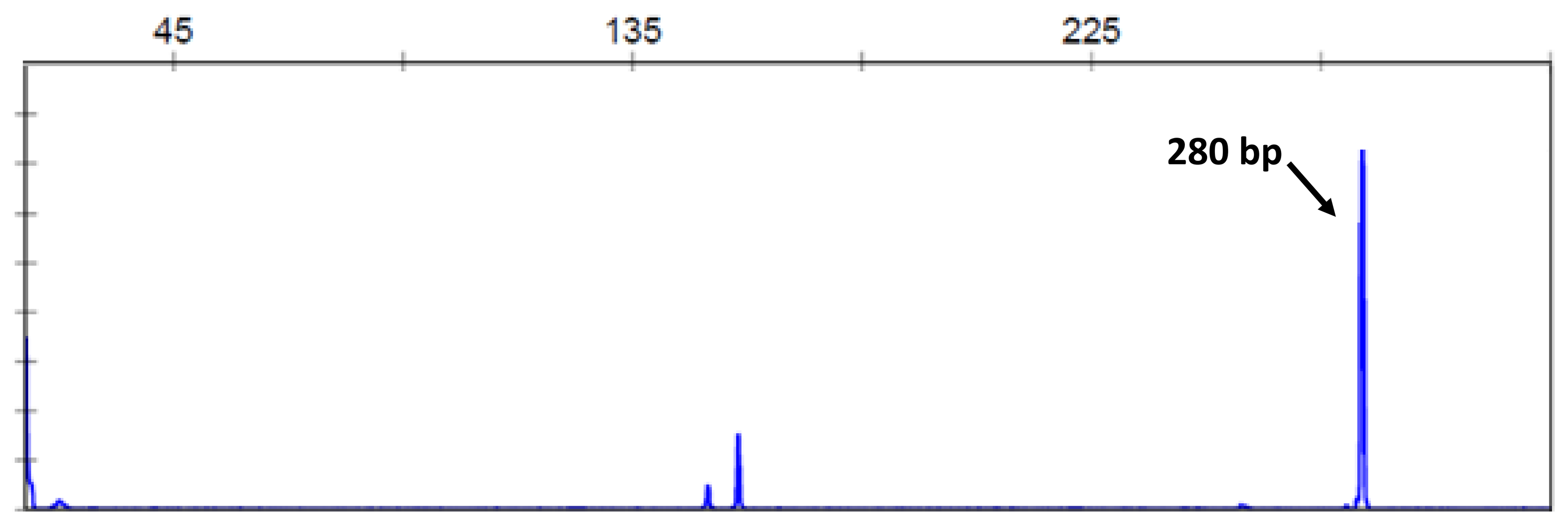 |
| 10-11 | 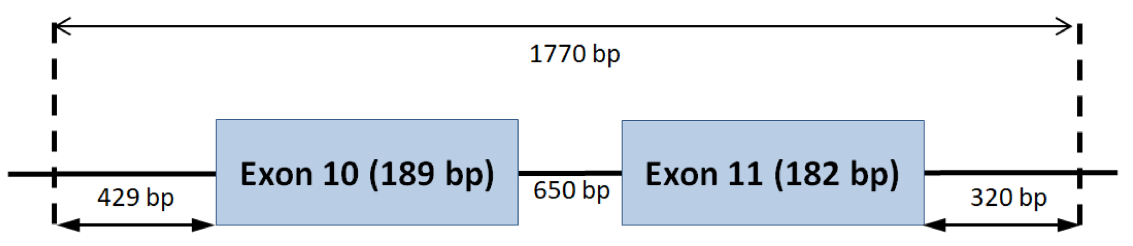 | 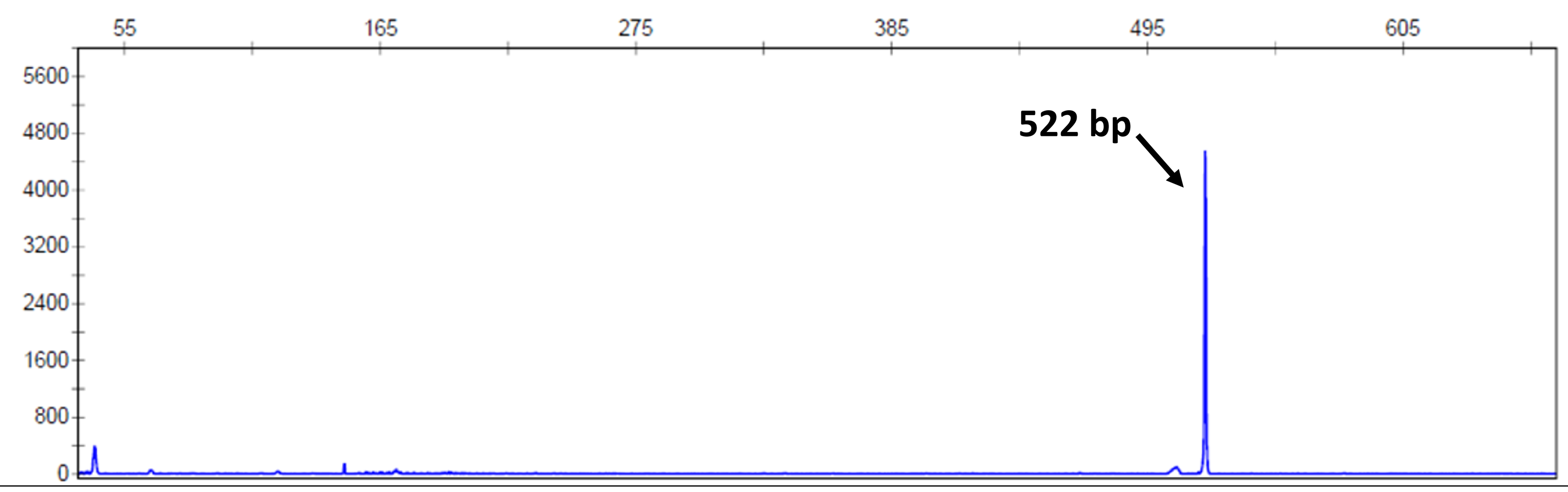 |
| 12 | 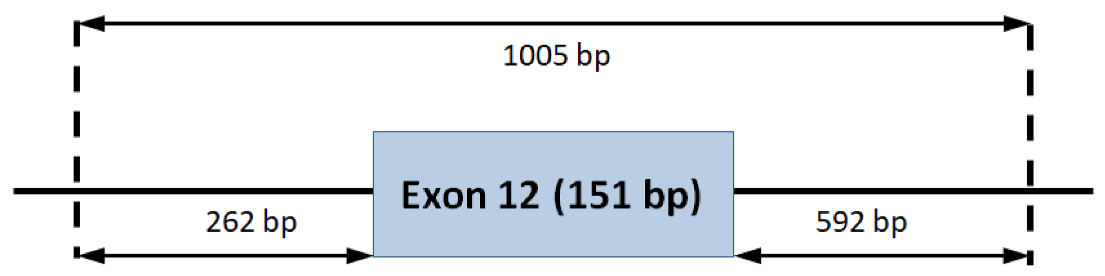 | 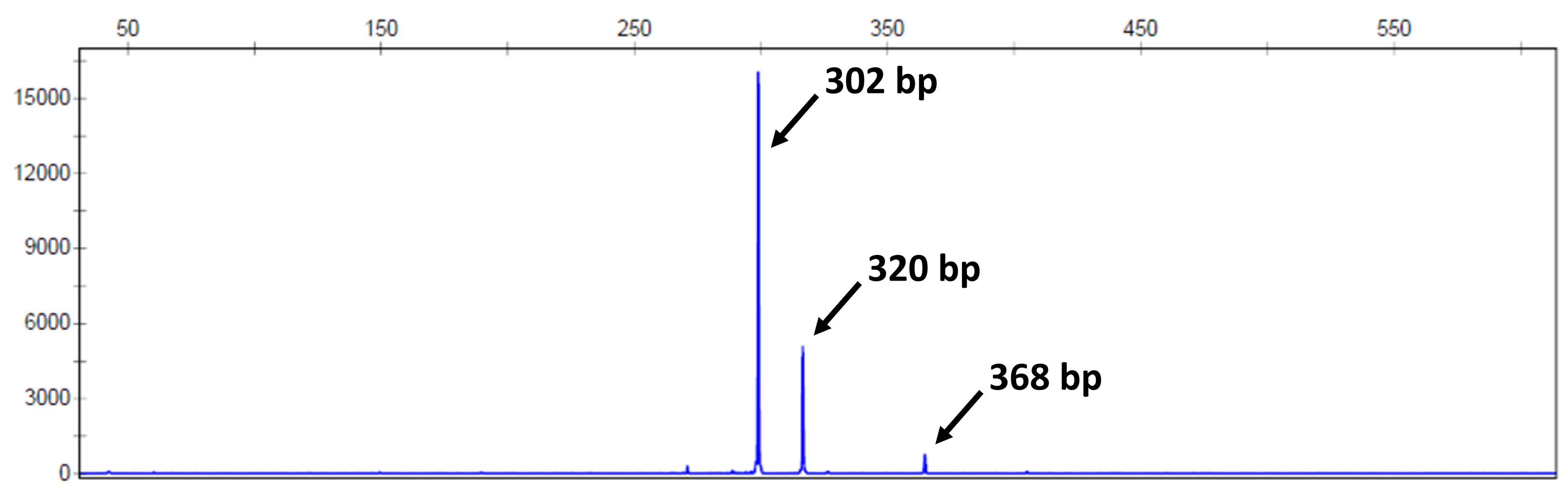 |
| 13 | 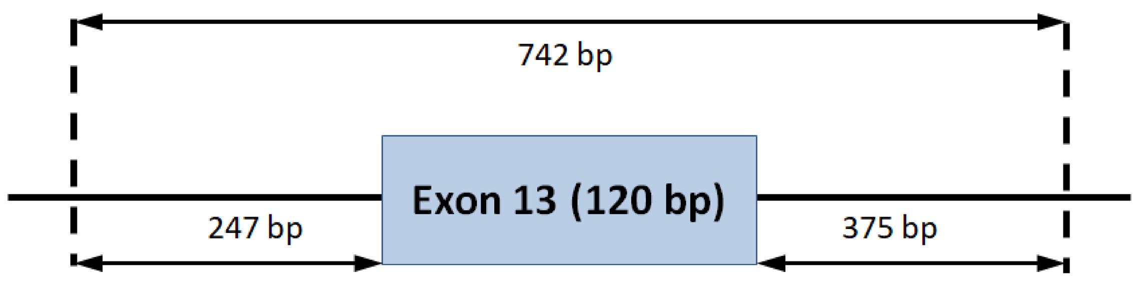 | 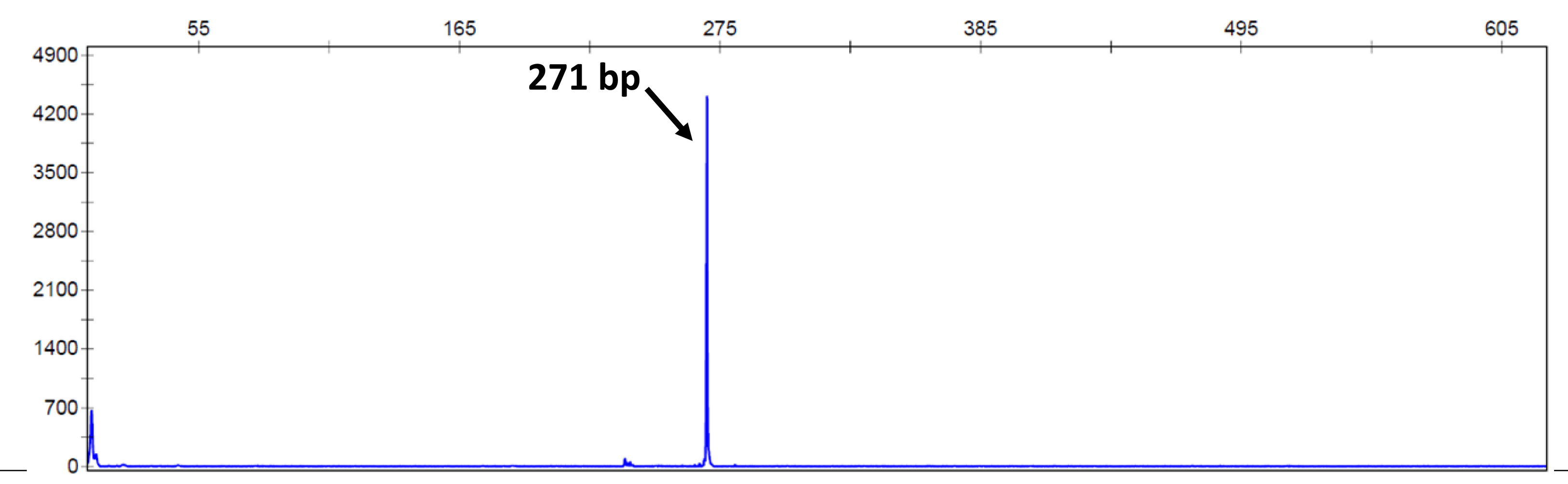 |
| 16 | 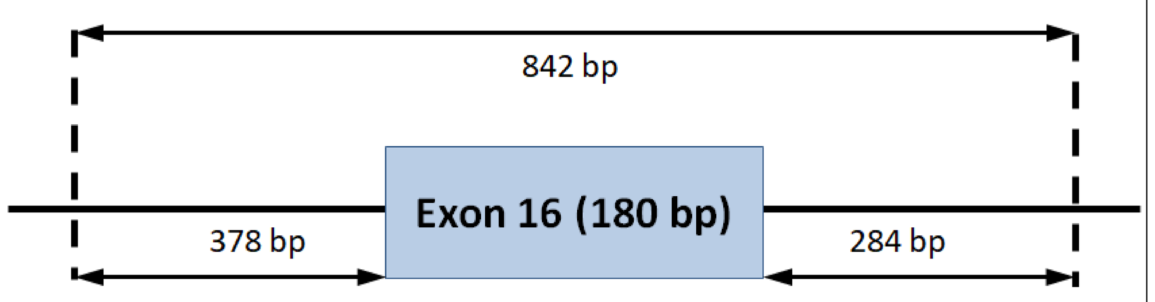 | 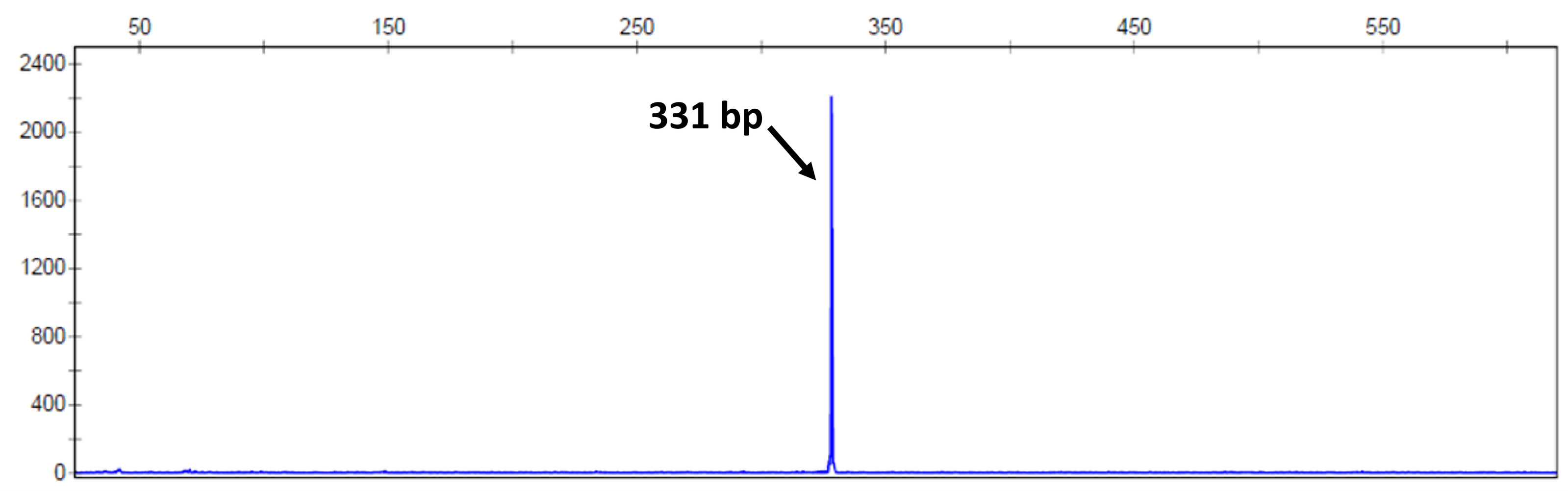 |
| 17 | 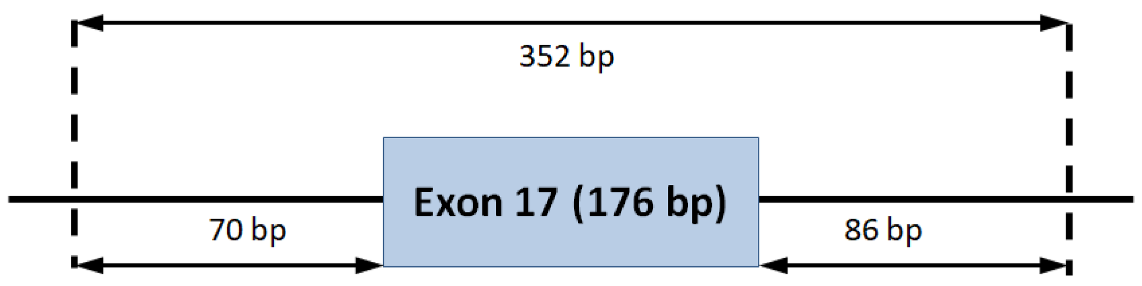 | 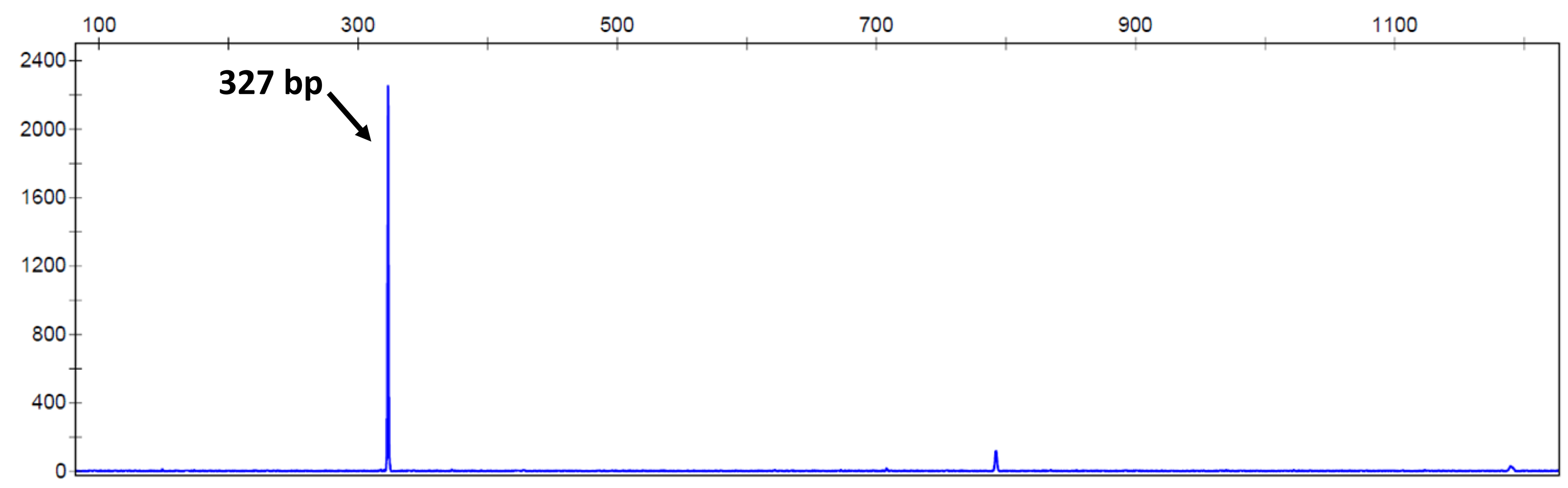 |

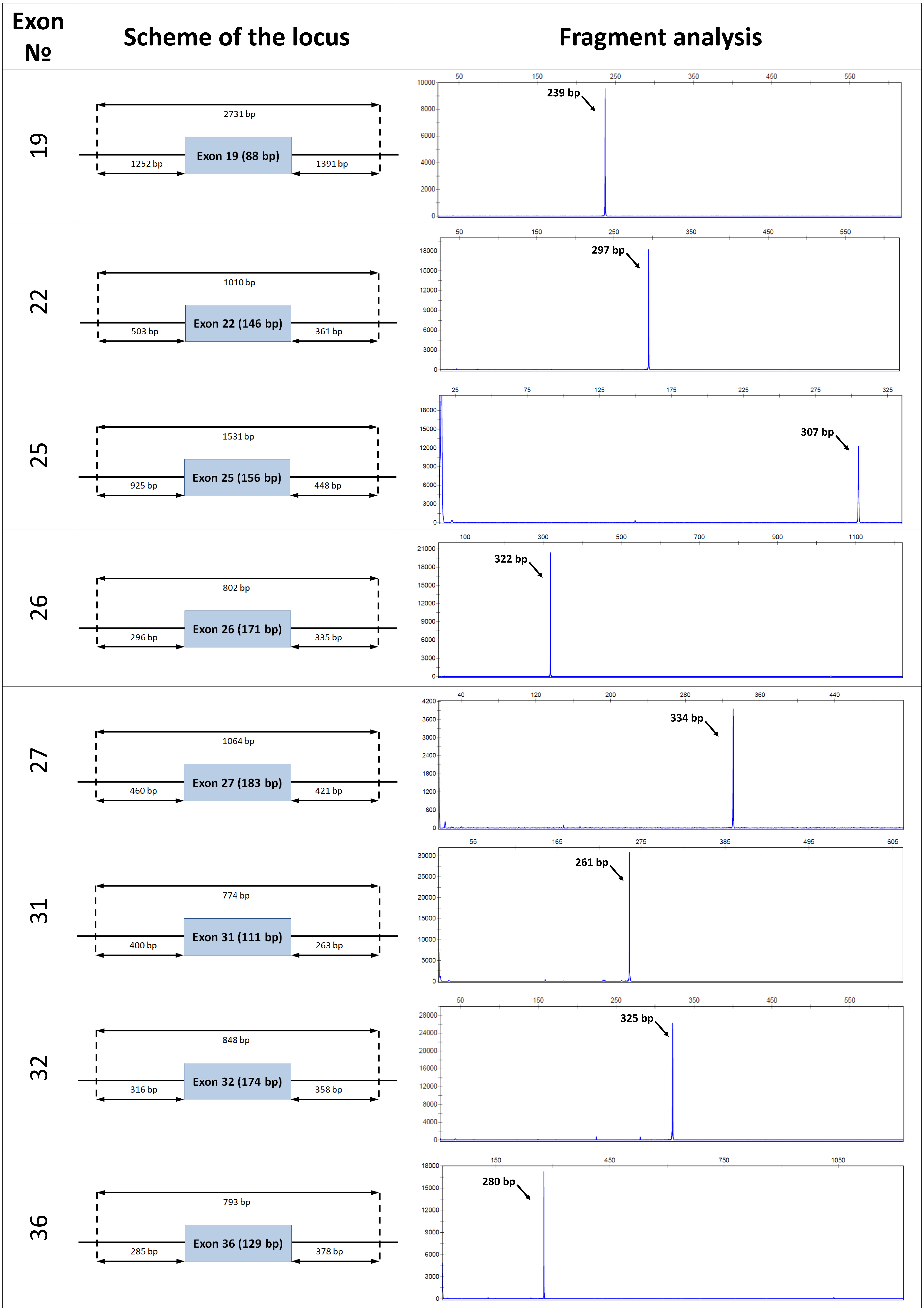

Exon №
Scheme of the locus
Fragment analysis
19
2731 bp
1252 bp
Exon 19 (88 bp)
1391 bp
239 bp
22
1010 bp
503 bp
Exon 22 (146 bp)
361 bp
297 bp
25
1531 bp
925 bp
Exon 25 (156 bp)
448 bp
307 bp
26
802 bp
296 bp
Exon 26 (171 bp)
335 bp
322 bp
27
1064 bp
460 bp
Exon 27 (183 bp)
421 bp
334 bp
31
774 bp
400 bp
Exon 31 (111 bp)
263 bp
261 bp
32
848 bp
316 bp
Exon 32 (174 bp)
358 bp
325 bp
36
793 bp
285 bp
Exon 36 (129 bp)
378 bp
280 bp

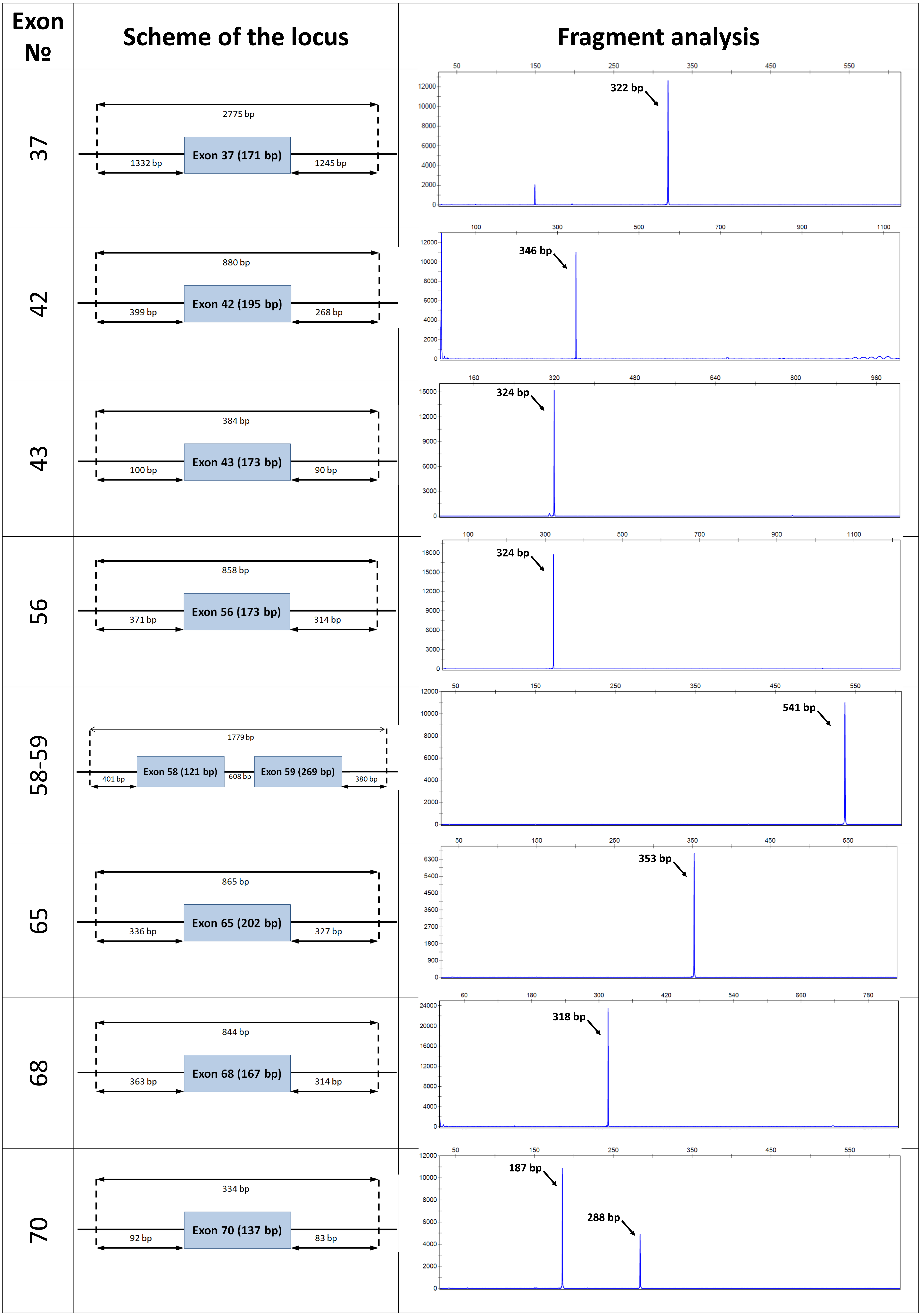

Exon №
Scheme of the locus
Fragment analysis
37
2775 bp
1332 bp
Exon 37 (171 bp)
1245 bp
322 bp
42
880 bp
399 bp
Exon 42 (195 bp)
268 bp
346 bp
43
384 bp
100 bp
Exon 43 (173 bp)
90 bp
324 bp
56
858 bp
371 bp
Exon 56 (173 bp)
314 bp
324 bp
58-59
1779 bp
401 bp
Exon 58 (121 bp)
608 bp
Exon 59 (269 bp)
380 bp
541 bp
65
865 bp
336 bp
Exon 65 (202 bp)
327 bp
353 bp
68
844 bp
363 bp
Exon 68 (167 bp)
314 bp
318 bp
70
334 bp
92 bp
Exon 70 (137 bp)
83 bp
187 bp
288 bp

| Exon № | Scheme of the locus | Fragment analysis |
| --- | --- | --- |
| 77 | 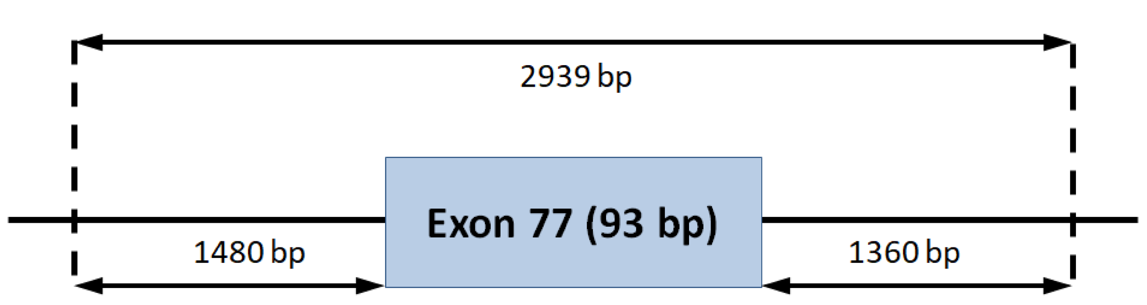  | 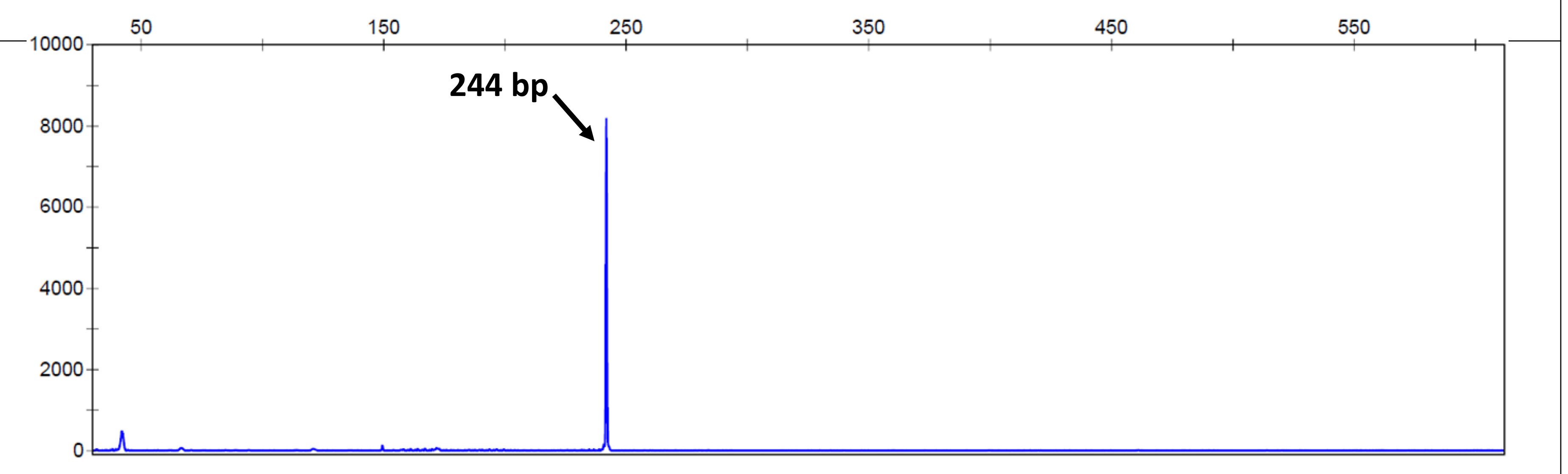  |

**Figure S1.** Design and functional validation of the wild-type minigene constructions. Left: genomic region contained the target *DMD* exon along with part of flanking introns were amplified and cloned into pSpl3-Flu2-TK vector. Right: fragment analysis by capillary electrophoresis displayed all products of the RT-PCR analysis. The identified transcript corresponds to a blue peak and their size is indicated by an asterisk.
