## Supplemental Figure S2 for "Revision of splicing variants in the *DMD* gene"

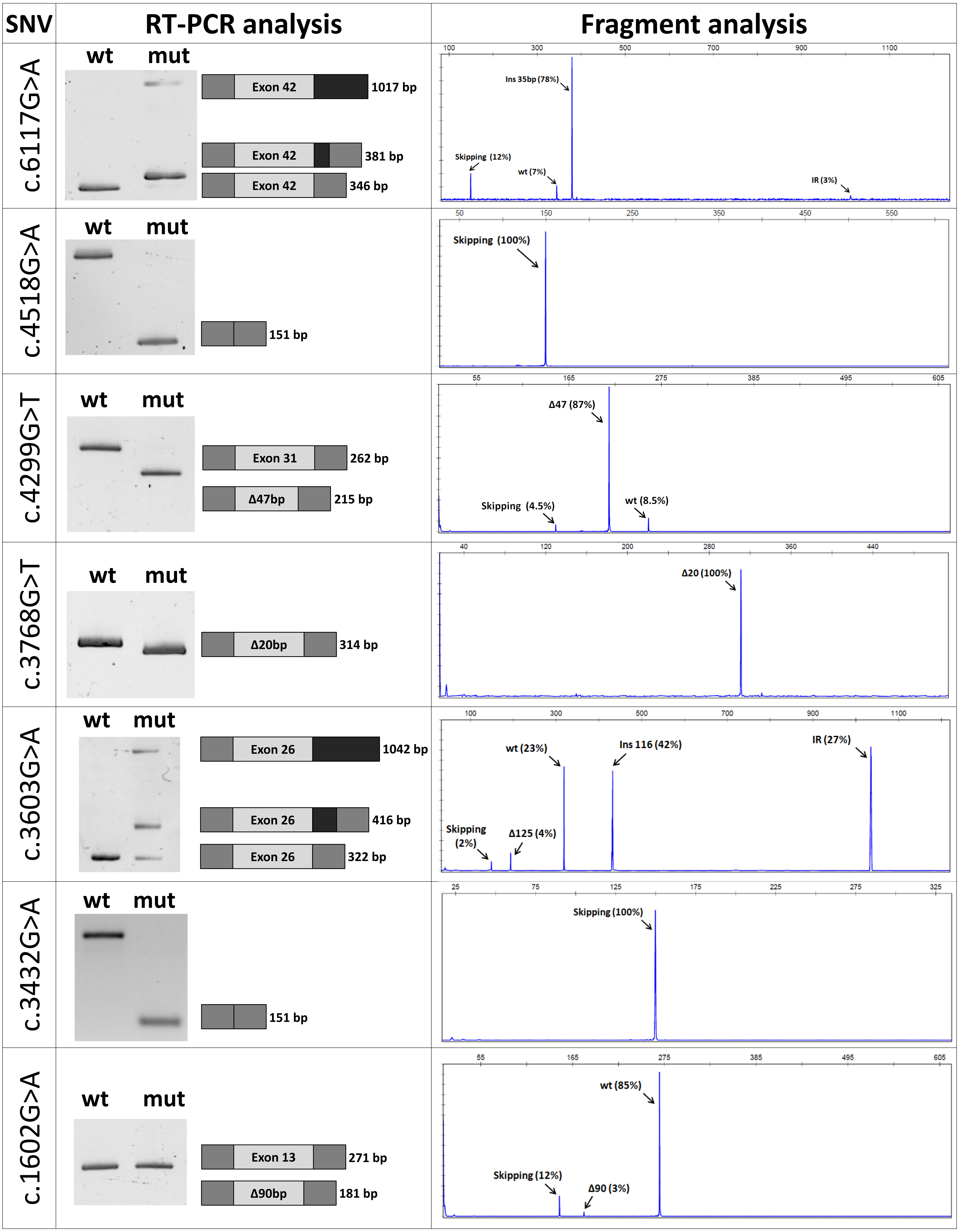

SNV
RT-PCR analysis
Fragment analysis
c.6117G>A
wt
mut
Exon 42
1017 bp
Exon 42
381 bp
Exon 42
346 bp
Ins 35bp (78%)
Skipping (12%)
wt (7%)
IR (3%)
c.4518G>A
wt
mut
151 bp
Skipping (100%)
c.4299G>T
wt
mut
Exon 31
262 bp
Δ47bp
215 bp
Δ47 (87%)
Skipping (4.5%)
wt (8.5%)
c.3768G>T
wt
mut
Δ20bp
314 bp
Δ20 (100%)
c.3603G>A
wt
mut
Exon 26
1042 bp
Exon 26
416 bp
Exon 26
322 bp
wt (23%)
Ins 116 (42%)
IR (27%)
Skipping (2%)
Δ125 (4%)
c.3432G>A
wt
mut
151 bp
Skipping (100%)
c.1602G>A
wt
mut
Exon 13
271 bp
Δ90bp
181 bp
wt (85%)
Skipping (12%)
Δ90 (3%)

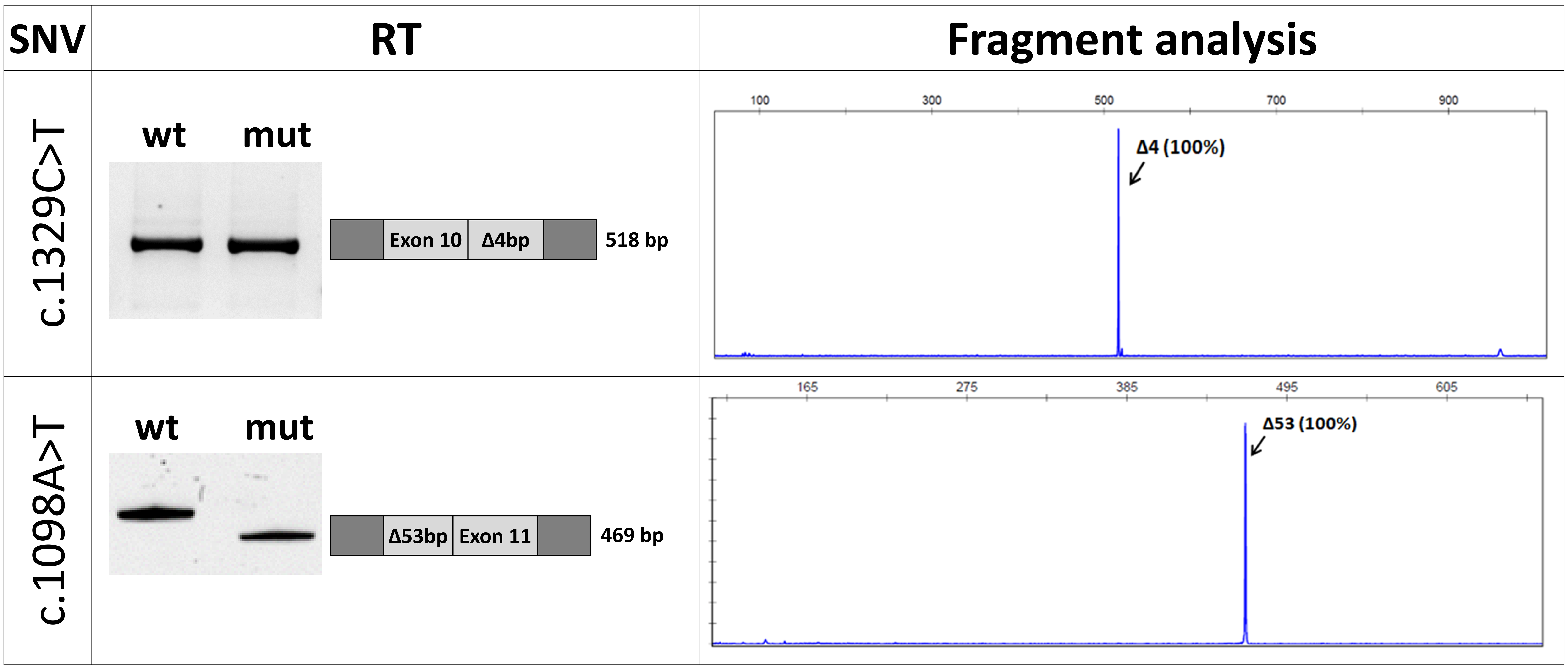


**Figure S2.** Functional analysis of synonymous variants in the *DMD* gene using the minigene expression system. Left: electrophoresis of RT-PCR products and schematic of the observed splicing changes. Right: fragment analysis of RT-PCR products for a variant-containing plasmid.

***wt*** *– wild type;* ***Δ*** *– deletion;* ***Ins*** *– insertion;* ***IR*** *– intron retention.*
