## Supplemental Figure S3 for "Revision of splicing variants in the *DMD* gene"

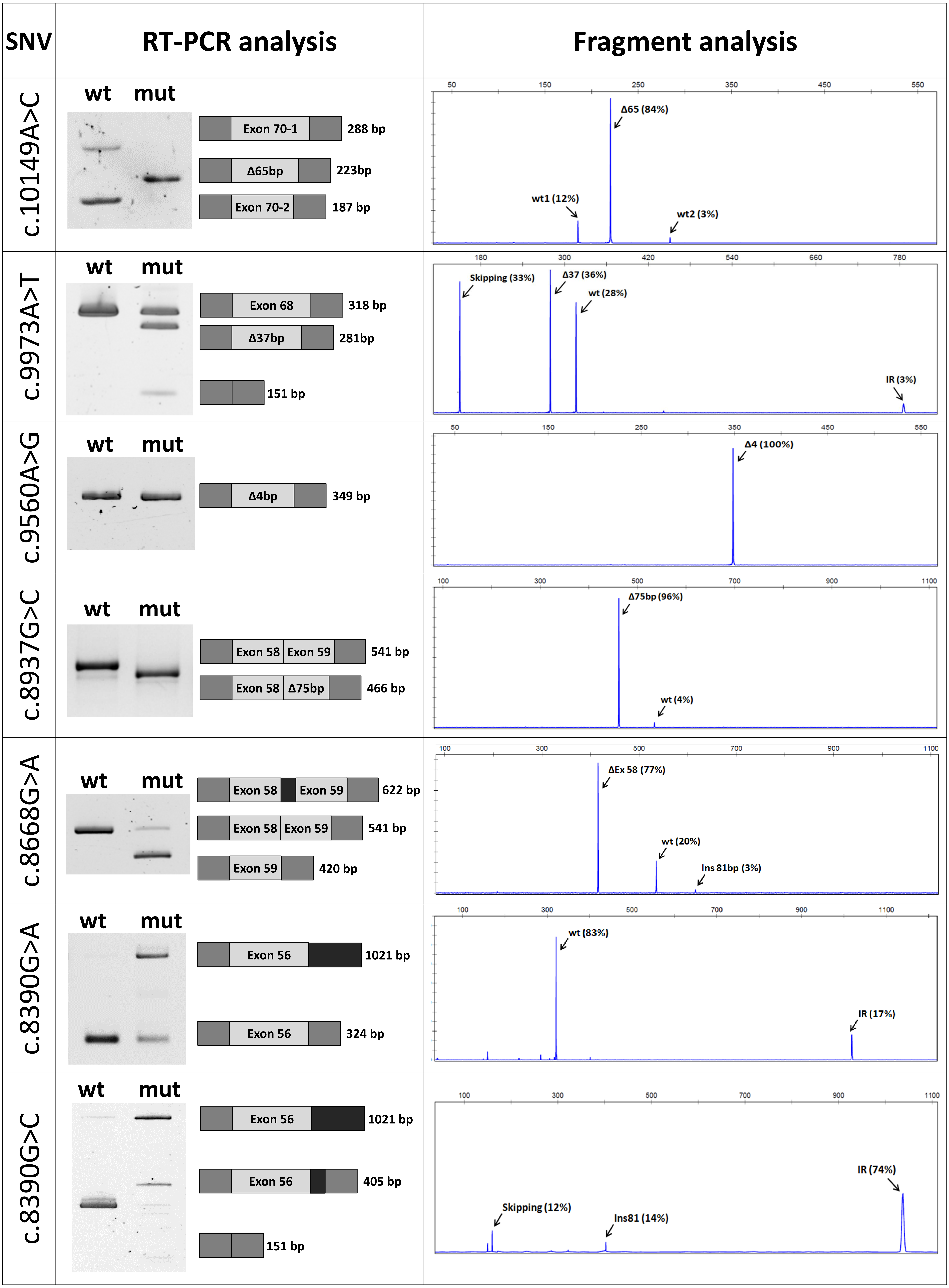

SNV
RT-PCR analysis
Fragment analysis
c.10149A>C
wt mut
Exon 70-1 288 bp
Δ65bp 223bp
Exon 70-2 187 bp
Δ65 (84%)
wt1 (12%)
wt2 (3%)
c.9973A>T
wt mut
Exon 68 318 bp
Δ37bp 281bp
151 bp
Skipping (33%)
Δ37 (36%)
wt (28%)
IR (3%)
c.9560A>G
wt mut
Δ4bp 349 bp
Δ4 (100%)
c.8937G>C
wt mut
Exon 58 Exon 59 541 bp
Exon 58 Δ75bp 466 bp
Δ75bp (96%)
wt (4%)
c.8668G>A
wt mut
Exon 58 Exon 59 622 bp
Exon 58 Exon 59 541 bp
Exon 59 420 bp
ΔEx 58 (77%)
wt (20%)
Ins 81bp (3%)
c.8390G>A
wt mut
Exon 56 1021 bp
Exon 56 324 bp
wt (83%)
IR (17%)
c.8390G>C
wt mut
Exon 56 1021 bp
Exon 56 405 bp
151 bp
Skipping (12%)
Ins81 (14%)
IR (74%)

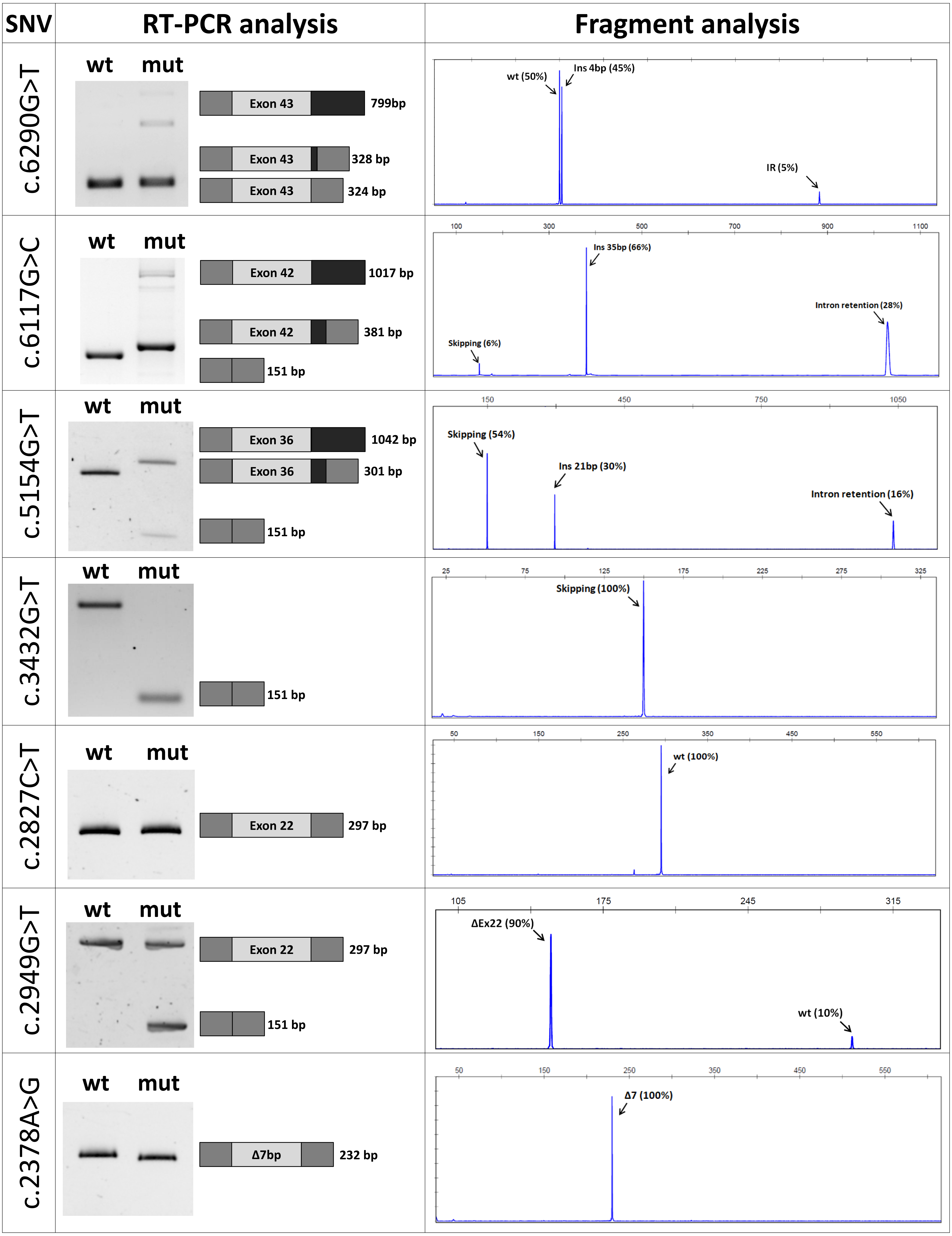
SNV
RT-PCR analysis
Fragment analysis
c.6290G>T
wt
mut
Exon 43
799bp
Exon 43
328 bp
Exon 43
324 bp
wt (50%)
Ins 4bp (45%)
IR (5%)
c.6117G>C
wt
mut
Exon 42
1017 bp
Exon 42
381 bp
151 bp
100
300
500
700
900
1100
Ins 35bp (66%)
Intron retention (28%)
Skipping (6%)
c.5154G>T
wt
mut
Exon 36
1042 bp
Exon 36
301 bp
151 bp
150
450
750
1050
Skipping (54%)
Ins 21bp (30%)
Intron retention (16%)
c.3432G>T
wt
mut
151 bp
25
75
125
175
225
275
325
Skipping (100%)
c.2827C>T
wt
mut
Exon 22
297 bp
50
150
250
350
450
550
wt (100%)
c.2949G>T
wt
mut
Exon 22
297 bp
151 bp
105
175
245
315
ΔEx22 (90%)
wt (10%)
c.2378A>G
wt
mut
Δ7bp
232 bp
50
150
250
350
450
550
Δ7 (100%)

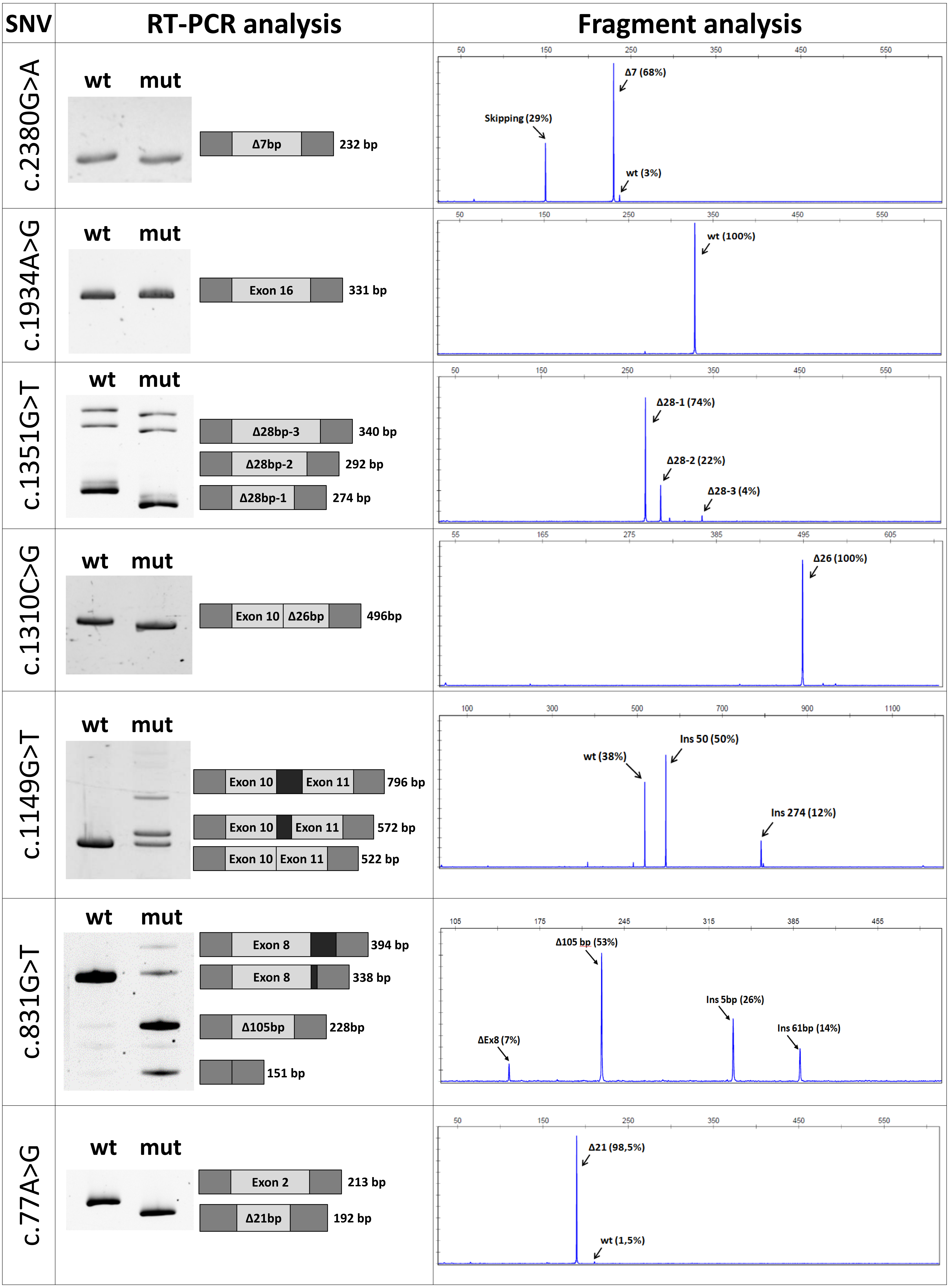


***wt*** *– wild type;* ***Δ*** *– deletion;* ***Ins*** *– insertion;* ***IR*** *– intron retention.*

**Figure S3.** Functional analysis of missense variants in the *DMD* gene using the minigene expression system. Left: electrophoresis of RT-PCR products and schematic of the observed splicing changes. Right: fragment analysis of RT-PCR products for a variant-containing plasmid.
