## Supplemental Figure S4 for "Revision of splicing variants in the *DMD* gene"

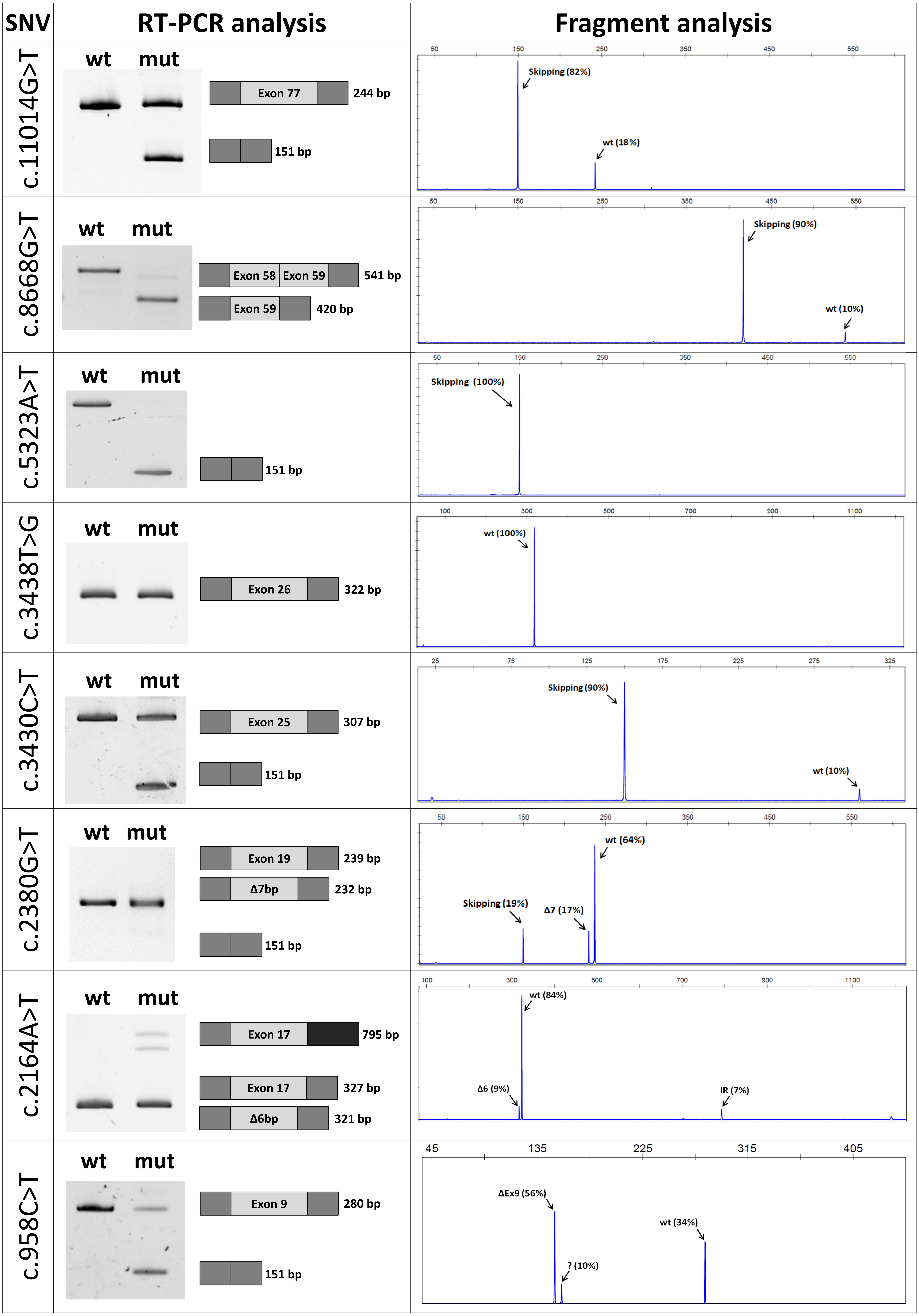


***wt*** *– wild type;* ***Δ*** *– deletion;* ***Ins*** *– insertion;* ***IR*** *– intron retention.*

**Figure S4.** Functional analysis of nonsense variants in the *DMD* gene the using minigene expression system. Left: electrophoresis of RT-PCR products and schematic of the observed splicing changes. Right: fragment analysis of RT-PCR products for a variant-containing plasmid*.*
