## Supplemental Table S1 for "Revision of splicing variants in the *DMD* gene"

### Table S3. Primers for cloning and mutagenesis

#### Primers for cloning

| Minigene construction | Primer name | Primer sequence 5'->3' |
|---|---|---|
| Exon 2 | XhoI-DMD Ex2F | AAAACTCGAGAACGACTATGGGTTTGGTCATA |
| | BamHl-DMD Ex2R | AAAAGGATCCCTTCTTCTGCTGGGTGACATA |
| Exon 8 | XhoI-DMD Ex8F | AAAACTCGAG CAGTGCACCATTTGAATTGACA |
| | BamHl-DMD Ex8R | AAAAGGATCCTTGCCTTTCATTTCATTGTGTGT |
| Exon 9 | BamHl-DMD Ex9R | AAAAGGATCCACCATTAGCAGCCTTTGTCAAA |
| | XhoI-DMD Ex9F | AAAACTCGAGTGCCGGATTGAAGAGTACCAT |
| Exons 10-11 | XhoI-DMD Ex10F | AAAACTCGAGTTCATTACCTAGGGCCTCTCA |
| | BamHl-DMD Ex11R | AAAAGGATCCTGCATGCTTCTGTGCCTTTTAT |
| Exon 12 | SacI-DMD Ex12F | AAAAGAGCTCCTTTGCACTTCAGTTAGCATGT |
| | BamHl-DMD Ex12R | AAAAGGATCCAAATCACATACCCTGCGTTGTT |
| Exon 13 | XhoI-DMD Ex13F | AAAACTCGAGGAGAACATCCTGCTGTACCTT |
| | BamHl-DMD Ex13R | AAAAGGATCCAGCTAATGATGAAATGGGTGCT |
| Exon 16 | XhoI-DMD Ex16F | AAAACTCGAGCCGTCAGGAATTAAGGGGAAA |
| | BamHl-DMD Ex16R | AAAAGGATCCAGTTCCCAGCATTTTGTTCGC |
| Exon 17 | XhoI-DMD Ex17F | AAAACTCGAGCCACCCCTTATTTCCTACAGT |
| | BamHl-DMD Ex17R | AAAAGGATCCCTAGGGACTCAACAAGCAGG |
| | XhoI-DMD Ex17-2F | AAAACTCGAGTGCCACTCCAAGCAGTCTTTA |
| | BamHl-DMD Ex17-2R | AAAAGGATCCTGCTGAAGTGAAAACACCACC |
| Exon 19 | XhoI-DMD Ex19F | AAAACTCGAGTGTAGCAAGTCATGAAAATGGC |
| | BamHl-DMD Ex19R | AAAAGGATCCACCACATCCCATTTTCTTCCAA |
| Exon 22 | XhoI-DMD Ex22F | AAAACTCGAGGCGTACATGGGTGTTCTTTCA |
| | BamHl-DMD Ex22R | AAAAGGATCCACCCACCAGTTTGAGAATGTG |
| Exon 25 | XhoI-DMD Ex25F | AAAACTCGAGACCTTCACACCCATTACACTG |
| | BamHl-DMD Ex25R | AAAAGGATCCACAGTTGTGTTCAAGAAAGTCAT |
| | XhoI-DMD Ex25-minF | AAAACTCGAGATGCCATCAGTCCCAATTTTAC |
| | BamHl-DMD Ex25-minR | AAAAGGATCCAAAGTAACGGTGAAGGGAGAC |
| | XhoI-DMD Ex25-maxF | AAAACTCGAGTCAGTGTTTTACAGCTGGGGT |
| | BamHl-DMD Ex25-maxR2 | AAAAGGATCCAGAGGCCAAACTTGGAAATGTT |
| Exon 26 | XhoI-DMD Ex26F | AAAACTCGAGAGCCTTGTTTTTCTCCATTCAC |
| | BamHl-DMD Ex26R | AAAAGGATCCTTTGCACAGTTTCAACGTACATT |
| Exon 27 | XhoI-DMD Ex27F | AAAACTCGAGAGGGGACCTTTCTATGTTAAGT |
| | BamHl-DMD Ex27R | AAAAGGATCCACAGAATGCTTTTCCCCAAAGA |
| Exon 31 | XhoI-DMD Ex31F | AAAACTCGAGTTGGTCCGATTTGAGTGTGAC |
| | BamHl-DMD Ex31R | AAAAGGATCCTCCATTCATCCAACACAGGAAA |
| Exon 32 | XhoI-DMD Ex32F | AAAACTCGAGGCAAGATTGGATTTGAGGACAT |
| | BamHl-DMD Ex32R | AAAAGGATCCACCTGGCATCCATTATTTCTGA |
| Exon 36 | XhoI-DMD Ex36F | AAAACTCGAGTTACACCCTTCTCTGTCACGA |
| | BamHl-DMD Ex36R | AAAAGGATCCGGTGTTCATTCAGATGGGCTA |

| | | |
|---|---|---|
| Exon 37 | XhoI-DMD Ex37F | AAAACTCGAGATGCATCAAACTTAAGGACGGT |
| | BamHl-DMD Ex37R | AAAAGGATCCGGTGACCCCTGAAAGTACGT |
| | XhoI-DMD Ex37-minF | AAAACTCGAGCTCGCTCTGTTTGGCTCTCT |
| | BamHl-DMD Ex37-minR | AAAAGGATCCGGTCTACTTGCCCCTTTCAG |
| | XhoI-DMD Ex37-maxF | AAAACTCGAGTTCTGCCTTTGTCAATGGTGG |
| | BamHl-DMD Ex37-maxR | AAAAGGATCCGGGAAGGTGAATTTTGAAGGG |
| Exon 42 | XhoI-DMD Ex42F | AAAACTCGAGCCCTCACCCAATCCCATATC |
| | BamHl-DMD Ex42R | AAAAGGATCCTACTGGTGGTTCCTTTCTGTC |
| Exon 43 | DMD-Ex43F | ATCAGGATGAGAAGGATCCAG |
| | NdeI -DMD Ex43R | AAAACATATGAATGCCCCAATCTGATTTACGA |
| | XhoI-DMD-Ex43-2F | AAAACTCGAGCACCATTTGCTACCTTTGGGA |
| | BamHl-DMD-Ex43-2R | AAAAGGATCCTTTTCCATGGAGGGTACTGAAA |
| Exon 56 | XhoI-DMD Ex56F | AAAACTCGAGGCACTGGGGTACACTTTATCA |
| | BamHl-DMD Ex56R | AAAAGGATCCAACCCTGTGATATGCCAAGAG |
| Exons 58-59 | SacI-DMD Ex58F | AAAAGAGCTCAGGCTTCCCATTATGTTCTGG |
| | NdeI-DMD Ex59R | AAAACATATGACTCCTGGCTTTTGATTAAGTG |
| Exon 65 | XhoI-DMD Ex65F | AAAACTCGAGTCAGGTGACAATTGACTCTGTT |
| | BamHl-DMD Ex65R | AAAAGGATCCACCCCTGAAGTGCATGGAAG |
| Exon 68 | XhoI-DMD Ex68F | AAAACTCGAGACCTGCATTTCACTTCTGGCT |
| | BamHl-DMD Ex68R | AAAAGGATCCGAAGGAGGAGGTACAGTTCAT |
| Exon 70 | XhoI-DMD Ex70F | AAAACTCGAGAGTGGGTGAAGCATGCTATCA |
| | BamHl-DMD Ex70R | AAAAGGATCCAGTGACTAGATATGACCACACA |
| | XhoI-DMD Ex70-2F | AAAACTCGAGCAAAACAAGTGTCATGGGGCA |
| | BamHl-DMD Ex70-2R | AAAAGGATCCTTTGGGAGTGAAGGAGGGTG |
| Exon 77 | XhoI-DMD Ex77F | AAAACTCGAGACTGTATGGATTTCTTCTTCCCT |
| | BamHl-DMD Ex77R | AAAAGGATCCACTTGACAAATAGAGGGTGGTA |

### Primers for mutagenesis

| Variant | Primer name | Primer sequence 5'->3' |
|---|---|---|
| c.77A>G | 77AG_R | CATTCTTACCTTAGAAAATTGTGCACTTACCCATTTTGTGAATGT |
| c.831G>T | 831GT_F | CAAATGCACTATTCTCAACATGTAAAGTGTGTAAAGGACAGC |
| c.958C>T | 958CT_F | CGGAGCCCATTTCCTTCATAGGTCTGTCAACATTTACTCT |
| c.1098A>T | 1098AT_F | GACACATTGCAAGCACAAGGTGAGATTTCTAATGATGTGGAAG |
| c.1149G>T | 1149GT_R | GTAAATTAACGTTTTAGTTTACATCATGAGTATGAAACTGGTCT |
| c.1310C>G | 1310CG_F | GGAATGCCTCAGGGTAGGTAGCATGGAAAAACAAAGCAAG |
| c.1351G>T | 1351GT_F | CAGTTTACATAGAGTTTTAATGTATCTCCAGAATCAGAAACTG |
| c.1934A>G | 1934AG_R | ACACCGGGCAAAGTTACCCAGCCATGCTTCCGT |
| c.2164A>T | 2164AT_F | CTGTGGATTCTGAAATTAGGTAAAGGTGAGAGCATCTTA |
| c.2378A>G | 2378AG_R | CTAAATCAACTCGTGTAATTACCACTCACCATCTGTTCCACCAG |
| c.2380G>A | 2380GA_R | CTAAATCAACTCGTGTAATTACTATTCACCATCTGTTCCACCAG |
| c.2380G>T | 2380GT_R | CTAAATCAACTCGTGTAATTACAATTCACCATCTGTTCCACCAG |
| c.2827C>T | 2827CT_F | GACACTTTGCCACCAATGTGCTATCAGGAGACCATGAGTGC |
| c.2949G>T | 2949GT_R | TTTGACATTCAAATATTCACAGACATGCAATTCCCCGAGTCTCTGct |

| | | |
|---|---|---|
| c.3438T>G | 3438TG_F | TGTTTTGTGGAAGGTCTAGGCCAGAAAGGAGGCCTTGAAG |
| c.3603G>A | 3603GA_F | GCAGTTGAAGAGATGAAAGTAAAAAAAAAAAAAGAAAAACTAAG |
| c.3603G>A | 3603GA_R | TTTACTTTCATCTCTTCAACTGCTTTCTGTAATTCA |
| c.4518G>A | 4518GA_F | TCACAGCTAAATCATTGTGTAGTATGTATTTCTGGTGGCAA |
| c.5323A>T | 5323AT_R | CTTGGAGTAGATCTTCCTACCTATCCAGTCTTAATTCTGTGTGAA |
| c.6117G>A | 6117GA_R | AATGAAAGTGCTTTGGTTTTACTTTCAGAGACTCCTCTTGCTTA |
| c.6290G>T | 6290GT_R | CAAGAAAAATATATGTGTTACCTACACTTGTCGGTCCTTGTACATtt |
| c.8390G>A | 8390GA_F | GGAAAAAGTCTCTCAACATTAAGTAGGAAAAGATGTGGAGCAA |
| c.8668G>A | 8668GA_R | ATAGTTCCACATTCAATTACTTCTGGGCTCCTGGTAGAGT |
| c.8668G>T | 8668GT_R | ATAGTTCCACATTCAATTACATCTGGGCTCCTGGTAGAGT |
| c.8937G>C | 8937GC_F | AGATCACCTCGAGAAAGTCAACGTACCGTCTACTTCTTTGCT |
| c.9560A>G | 9560AG_F | GGCTGCTGAATGTTTATGGTACGTACGTATGGCATGTTTTTA |
| c.10149A>C | 10149AC_F | CTAAAAAACAAATTTCGAACCAACAGGTATTTTGCGAAGCATCC |
| c.11014G>T | 11014GT_F | CAACTCCTTCCCTAGTTCAAGATGTAAGCTCCAATACCTAGAA |
| c.3430C>T | 3430CTF | GGATCACATGTGCCAATAGGTATAGACAATCTCTTTCACT |
| c.3432G>A | 3432GAF | GGATCACATGTGCCAAcAAGTATAGACAATCTCTTTCACT |
| c.3432G>T | 3432GTF | GGATCACATGTGCCAAcATGTATAGACAATCTCTTTCACT |
| c.9973A>T | 9973ATR | TTTTGGTTCCTAATACCAGAATCCAATGATTGGACACTC |
| c.6117G>C | 6117GCR | AAGTGCTTTGGTTTTACGTTCAGAGACTCCTCTTGC |
| c.8390G>C | 8390GCF | AAAGTCTCTCAACATTACGTAGGAAAAGATGTGGAGC |
| c.1329C>T | 1329CTR | ACAAATAAGGACTTACTTACTTTGTTTTTCCATGCT |
| c.1602G>A | 1602 MUT-F | /Pi/-ACAACTTAAAGTCAGATTATTTGCTTAGT |
| | 1602 MUT-R | /Pi/-TCTTCCAAAGCAGCAGTTGCG |
| c.3768G>T | 3768 MUT-F | /Pi/-GCTGAATGGTAAATGCAAGACTTTGG |
| | 3768 MUT-R | /Pi/-CTAGTGCAGAGCCACTGGTAGTTG |
| c.4299G>T | 4299 MUT-F | /Pi/-AACATAATCAGGGTAAGGAGGCTG |
| | 4299 MUT-R | /Pi/-TCTTCATTTCTTCTAAACTGATCTCATGACT |
| c.5154G>T | 5154 MUT-F | /Pi/-AAGACGTGCTTAATGTAGCAAATAAAAT |
| | 5154 MUT-R | /Pi/-CTTTTTGCTGGGGTTTCTTTTTCTCT |
