## Supplemental Table S3 for "Revision of splicing variants in the *DMD* gene"

| Variant | | PMID | Phenotype | Exon number | SpliceAI prediction | Expected splicing events | Observed splicing events | | | Clinical features (if any) |
|---|---|---|---|---|---|---|---|---|---|---|
| cDNA | Protein | | | | | | % in-frame isoforms | % out-of-frame isoforms | % WT transcript | |
| c.6117G>A | Lys2039= | 20485447 | BMD | 42 | DL: 0.72;<br>DG: 0.39 | Exon 42 skipping (in-frame);<br>Cryptic intronic splice site activation | Exon skipping (p.His1975_Lys2039del) — 12% | Partial intron retention, insertion of 35 nt (p.N2040Vfs*14) — 81% | 7% | - |
| c.4518G>A | Val1506= | 19937601 | IMD | 32 | DL: 0.62;<br>DG: 0.54 | Exon 32 skipping (in-frame) | Exon skipping (p.Lys1449_Val1506del) — 100% | - | - | - |
| c.4299G>T | G1443= | 32194622 | BMD | 31 | DG: 0.48;<br>DL: 0.59 | New donor splice site;<br>Exon 31 skipping (in-frame) | Exon skipping (p.Ile1413_Lys1449del) — 4.5% | Exon truncation, deletion of 47 nt (p.Gly1433Glufs*20) — 87% | 8,50% | - |
| c.3768G>T | G1256= | 32194622 | BMD | 27 | DG: 0.99;<br>DL: 0.74 | New donor splice site<br>Exon 27 skipping (in-frame) | - | Exon truncation, deletion of 20 nt (p.Lys1257Serfs*7) — 100% | - | - |
| c.3603G>A | Lys1201= | 19959795 | DMD | 26 | DL: 0.89 | Exon 26 skipping (in-frame) | Exon skipping (p.Val1160_Lys1201del) — 4% | Intron retention (p.R1202Vfs*25) — 71% | 23% | - |
| c.3432G>A | Gln1144= | 19783145 | BMD | 25 | DL: 0.98 | Exon 25 skipping (in-frame) | Exon skipping (p.Leu1093_Gln1144del) — 100% | - | - | Immunohistochemical labelling using antibodies DYS1,2,3: DYS1 and DYS2 normal, DYS3 decreased intensity |
| c.1602G>A | Lys534= | 31443951;<br>31706698 | BMD | 13 | DL: 0.91 | Exon 13 skipping (in-frame) | Exon skipping (p.Gln497_Leu536del) — 12%<br>Exon truncation, deletion of 90 nt (p.Arg506_Val535del) — 3% | - | 85% | The male proband presented at age 10 years with a persistently elevated serum creatine kinase (CK) of 19,372 U/L (normal levels < 200 U/L) and myalgia with exercise, but no associated weakness. He had a history of mild speech delay and was diagnosed with autism spectrum disorder at the age of 14 years and major depression at 16 years of age. At 18 years of age, he experiences muscle cramps when exercising for longer than one hour. His-power remains normal. He has mild tendoachilles and hamstring contractures. Electrocardiography and echocardiograms have been normal. Family history revealed that the maternal grandfather was diagnosed with Becker muscular dystrophy (with superimposed in-framelammatory myositis) after presenting at the age of 50 with mild limb-girdle muscle weakness and a modestly elevated CK. He was noted to have large calves and reported difficulty playing sport in childhood. Muscle biopsy showed patchy dystrophin staining and rimmed vacuoles. |
| c.1329C>T | Ser443= | 32559196 | BMD | 11 | DG: 0.98;<br>DL: 0.61 | New donor splice site<br>Exon 11 skipping (out-of-frame) | - | Exon truncation, deletion of 4 nt (p.Ser443Ilefs*6) — 100% | - | 23 yrs, Ambulant |
| c.1098A>T | Gly366= | 11524473 | DMD | 10 | DG: 0.98 | Cryptic exonic splice site activation | - | Exon truncation, deletion of 53 nt (p.Glu367Valfs*14) — 100% | - | - |
| c.10149A>C | p.(Lys3383Asn) | 19937601;<br>25220406 | B/DMD | 70 | AG: 0.84 | New acceptor splice site | - | Exon truncation, deletion of 65 nt (p.Thr3363Valfs*48) — 85% | 15% | Unknown phenotype |
| c.9973A>T | p.(Arg3325Trp) | 32169422 | Dystrophinopathy | 68 | DL: 0.42;<br>DG: 0.26 | Exon 68 skipping (out-of-frame);<br>New donor splice site | - | Exon skipping (p.Asn3314Thrfs*4) — 36%<br>Exon truncation, deletion of 37 nt (p.Ala3270Valfs*8) — 33%<br>Intron retention (p.Arg3327*) — 3% | 28% | - |
| c.9560A>G | p.(Asp3187Gly) | 17259292 | DMD | 65 | DG: 0.99;<br>DL: 0.70 | New donor splice site;<br>Exon 65 skipping (out-of-frame) | - | Exon truncation, deletion of 4 nt (p.Asp3187Glyfs*95) — 100% | - | - |
| c.8937G>C | p.(Lys2979Asn) | 27593222 | MD | 59 | DL: 0.79 | Exon 59 skipping (out-of-frame) | Exon truncation, deletion of 75 nt (p.Val2955_Lys2979del) — 96% | - | 4% | - |
| c.8668G>A | p.(Glu2890Lys) | 22776072;<br>25220406 | BMD | 58 | DL: 0.41 | Exon 58 skipping (out-of-frame) | - | Exon skipping (p.Ala2850Serfs*29) — 80% | 20% | - |
| c.8390G>A | p.(Arg2797Lys) | 22776072 | BMD | 56 | DL: 0.77 | Exon 56 skipping (out-of-frame) | - | Intron retention (p.Ser2798*) — 17% | 83% | - |
| c.8390G>C | p.(Arg2797Thr) | 11524473 | DMD | 56 | DL: 0.97 | Exon 56 skipping (out-of-frame) | - | Intron retention (p.Ser2798*) — 88%<br>Exon skipping (p.Asp2740Valfs*8) — 12% | - | - |
| c.6290G>T | p.(Gly2097Val) | 30907348 | DMD | 43 | DG: 0.61 | New donor splice site | - | Partial intron retention, insertion of 4 nt (p.Arg2098Valfs*4) — 45%<br>Intron retention p.(Arg2098*) — 5% | 50% | - |
| c.6117G>C | p.(Lys2039Asn) | 19937601;<br>25525159 | DMD | 42 | DL: 0.87;<br>DG: 0.45 | Exon 42 skipping (in-frame);<br>Cryptic intronic splice site activation | Exon skipping (p.His1975_Lys2039del) — 6% | Partial intron retention, insertion of 35 nt (p.N2040Vfs*14) — 66%<br>Intron retention (p.N2040Vfs*14) — 28% | - | - |
| c.5154G>T | p.(Lys1718Asn) | 32528171 | Muscular dystrophy, limb girdle | 36 | DL: 0.98;<br>DG: 0.31 | Exon 36 skipping (in-frame);<br>Cryptic intronic splice site activation | Exon skipping (p.Glu1676_Lys1718del) — 54% | Intron retention (p.R1719Vfs*6) — 46% | - | - |
| c.3432G>T | p.(Gln1144His) | 23756440;<br>29604111 | BMD | 25 | DL: 0.98 | Exon 25 skipping (in-frame) | Exon skipping (p.Leu1093_Gln1144del) — 100% | - | - | Disease onset: 8 |
| c.2827C>T | p.(Arg943Cys) | 25474345 | Stargardt macular dystrophy | 22 | AG: 0.27 | New acceptor splice site | - | - | 100% | - |
| c.2949G>T | p.(Gln983His) | 28859693;<br>29973226 | DMD | 22 | DL: 0.87 | Exon 22 skipping (out-of-frame) | - | Exon skipping (p.Ile935Serfs*30) — 90% | 10% | Dystrophin immunostaining - negative |
| c.2378A>G | p.(Asn793Ser) | 22776072;<br>25220406 | BMD | 19 | DG: 0.98;<br>DL: 0.71 | Cryptic exonic splice site activation;<br>Exon 19 skipping (out-of-frame) | - | Exon truncation, deletion of 7 nt (p.Val792Argfs*15) — 100% | - | - |
| c.2380G>A | p.(Glu794Lys) | 29365344 | DMD | 19 | DL: 0.89;<br>DG: 0.56 | Exon 19 skipping (out-of-frame);<br>Cryptic exonic splice site activation | - | Exon truncation, deletion of 7 nt (p.Val792Argfs*15) — 68%<br>Exon skipping (p.Ala765Argfs*15) — 29% | 3% | - |
| c.1934A>G | p.(Asp645Gly) | 7981690 | DMD | 16 | DG: 0.35 | New donor splice site | - | - | 100% | - |
| c.1351G>T | p.(Asp451Tyr) | 22776072;<br>32597815 | BMD | 12 | AG: 0.87 | New acceptor splice site | - | Exon truncation, deletion of 28 nt (p.Asn444Lysfs*5) — 100% | - | - |
| c.1310C>G | p.(Ala437Gly) | 19937601;<br>32559196 | DMD | 11 | DG: 0.96 | New donor splice site | - | Exon truncation, deletion of 26 nt (p.Val436Phefs*3) — 100% | - | 16 y.o., Non ambulant |
| c.1149G>T | p.(Glu383Asp) | 27122458 | B/DMD | 10 | DL: 0.72 | Exon 10 skipping (in-frame) | - | Partial intron retention, insertion of 50/274 nt (p.Gly384Valfs*3) — 62% | 38% | Unknown phenotype |
| c.831G>T | p.(Gln277His) | 30833962 | DMD | 8 | DL: 0.99 | Exon 8 skipping (out-of-frame) | Exon truncation, deletion of 105 nt (p.Val243_Gln277del) — 53% | Partial intron retention, insertion of 5 nt (p.I278Vfs*7) — 26%<br>Partial intron retention, insertion of 61 nt (p.I278Vfs*20) — 14%<br>Exon skipping (p.Val218Hisfs*9) — 7% | - | 2 y.o. CK (U/L): 14082; SCRN (µmol/L): 14 |
| c.77A>G | p.(Asn26Ser) | 30833962;<br>31081998 | DMD | 2 | DG: 0.92 | New donor splice site | Exon truncation, deletion of 21 nt (p.Val25_Lys31del) — 100% | - | - | 8 y.o., Expression levels of dystrophin C-terminus - Absence, CK (U/L): 11951 |
| c.11014G>T | p.(Gly3672*) | - | | 77 | DL: 0.46 | Exon 77 skipping (in-frame) | Exon skipping (p.Glu3642_Gly3672del) — 82% | - | 18% | - |
| c.8668G>T | p.(Glu2890*) | - | | 58 | DL: 0.6 | Exon 58 skipping (out-of-frame) | - | Exon skipping (p.Ala2850Serfs*29) — 90% | 10% | - |
| c.5323A>T | p.(Lys1775*) | - | | 37 | DL: 0.64 | Exon 37 skipping (in-frame) | Exon skipping (p.Arg1719_Lys1775del) — 100% | - | - | - |
| c.3438T>G | p.(Tyr1146*) | - | | 26 | AG: 0.78 | New acceptor splice site | - | - | 100% | - |
| c.3430C>T | p.(Gln1144*) | 19959795 | DMD | 25 | DL: 0.55 | Exon 25 skipping (in-frame) | Exon skipping (p.Leu1093_Gln1144del) — 90% | - | 10% | Skipping of the mutated exon observed at the RNA level |
| c.2380G>T | p.(Glu794*) | 11524473 | DMD | 19 | DL: 0.8;<br>DG: 0.25 | Exon 19 skipping (out-of-frame);<br>Cryptic exonic splice site activation | - | Exon skipping (p.Ala765Argfs*15) — 19%<br>Exon truncation, deletion of 7 nt (p.Val792Argfs*15) — 17% | 64% | - |
| c.2164A>T | p.(Lys722*) | - | | 17 | DG: 0.67 | New donor splice site | Exon truncation, deletion of 6 nt (p.Lys722_Arg723del) — 9% | Intron retention (p.Leu724*) — 7% | 84% | - |
| c.958C>T | p.(Gln320*) | - | | 9 | DL: 0.56 | Exon 9 skipping (in-frame) | Exon skipping (p.Ile278_Gln320del) — 56% | - | 34% | |
